# CCN1 forms a complex with GPC4 and heparin to fine-tune signaling activities for cortical neural stem cell maintenance

**DOI:** 10.1101/2025.05.16.654402

**Authors:** Jiangli Zheng, Shaojun Qi, Huan Wu, Jun Wu, Guo Chen, Huanhuan Joyce Wang, Han Pan, Ji Li, Cuiqin Cheng, Ke Tang, Lester F. Lau, Qin Shen

## Abstract

Radial glial cells (RGCs) are the neural stem cells (NSCs) in the developing brain, capable of self-renewing and generating neurons and glia in order. The developmental behaviors of NSCs are determined by extrinsic factors from the NSC niche, together with intrinsic factors. Here, we show that Cellular Communication Network 1 (CCN1), a secreted matricellular protein, is an autocrine factor derived from RGCs in the embryonic telencephalon. Loss of *Ccn1* in RGCs leads to accelerated lineage progression of RGCs, and causes premature cell cycle exit and neuronal differentiation, resulting in a reduction in NSC pool size, and in turn, at the expense of the perinatal gliogenesis. Mechanistically, CCN1 interacts with an RGC-expressed membrane protein GPC4, one of the heparan sulfate proteoglycans (HSPGs), and is required for GPC4 to maintain NSCs through the Sonic Hedgehog (Shh) signaling pathway, depending on the binding of heparin to CCN1. Our work provides a potential mechanism for the dynamic and sophisticated interconnections of niche components that meet the needs of context-dependent maintenance, differentiation, and fate determination of NSCs.

## Introduction

In mammals, neural stem cells (NSCs) first appear as neuroepithelial cells lining the ventricular surface. They transform into radial glial cells (RGCs) during embryonic development (Gotz and Huttner, 2005). RGCs divide symmetrically to self-renew, or they go through asymmetric division to produce neurons directly or indirectly via intermediate progenitor cells (IPCs). These newborn neurons migrate through the intermediate zone (IZ) and finally follow the inside-out fashion to populate their destined layer of the cortical plate (CP), which becomes the highly wired six-layered structure after birth. At the end of neurogenesis, RGCs generate astrocytes and oligodendrocytes (Loo et al., 2019; Molyneaux et al., 2007; Mukhtar and Taylor, 2018; Noctor et al., 2004).

Both embryonic and adult NSCs are located in a microenvironment, i.e., stem cell niche, composed of neighboring cells, soluble factors, and extracellular matrix (Bjornsson et al., 2015). Lines of evidence demonstrate that the maintenance and the sequential fate specification of NSCs are orchestrated by dynamic competition and cooperation between niche factor signaling pathways, such as Notch (Mizutani and Saito, 2005; Yoon and Gaiano, 2005), FGF2 (Inglis-Broadgate et al., 2005; Korada et al., 2002; Raballo et al., 2000; Vaccarino et al., 1999), Shh (Komada et al., 2008; Wang et al., 2016), and Wnt (Chenn and Walsh, 2003; Noles and Chenn, 2007; Woodhead et al., 2006; Zechner et al., 2003), as well as their downstream transcription factors (Gauthier-Fisher and Miller, 2013). Therefore, understanding the mechanisms of how the niche components fine-tune the availability dynamics of various ligands and control behaviors of NSCs has implications for NSC transplantation and regenerative medicine.

*Ccn1* (cellular communication network 1), *a.k.a. Cyr61* (cysteine-rich protein 61), has been discovered as an immediate-early gene activated in response to growth factor stimulation (O’Brien et al., 1990). It encodes one of the six family members of CCN proteins, which have been called for attention to their expression and function in the nervous system recently (Malik et al., 2015). CCN proteins are involved in multiple biological processes, including vasculogenesis, osteogenesis and chondrogenesis, nerve conduction, and muscular contraction (Perbal, 2004). Besides, CCN family members have four highly conserved domains, namely, the IGFBP (insulin-like growth factor-binding protein-like), the VWC (von Willebrand type C), the TSP-1 (thrombospondin type 1 repeat), and the CT (C-terminal) domains, which enable binding to multiple partners and guarantee the multifacet signaling modifications throughout life (Lau, 2011; Takigawa, 2017). Recently, CCN1 was identified in the ependymal cells of the adult V-SVZ (ventricular-subventricular zone) stem cell niche and restricted the adult NSC pool size through EGFR signaling (Wu et al., 2020). However, whether it has a similar role in the embryonic brain remains elusive.

In this study, we explored the expression pattern, functions, and underlying mechanisms of CCN1 during cortical development. We found that CCN1 is an RGC-derived autocrine matricellular factor. Conditional deletion of *Ccn1* in neural progenitor cells using the *Nestin*-Cre^+/-^/*Ccn1*^fl/fl^ (we designated as CcKO) mice disturbs NSC maintenance, accelerates RGC lineage progression, and induces premature consumptive neuronal differentiation at the expense of perinatal gliogenesis. We further discovered that CCN1 depends on heparin-binding and is required for RGC membrane-anchored GPC4 (Glypican-4) to maintain NSCs by regulating the Shh signaling pathway. These results suggest a context-dependent elaborate tuning of niche factor accessibility to NSCs through the GPC4-CCN1-heparin complex to regulate their fate.

## Results

### *Ccn1* is an autocrine matricellular niche factor of radial glial cells during cortical development

CCN1 has been identified in the conditioned medium of RGC culture (Siqueira et al., 2018). To investigate the origin of CCN1 in the developing forebrain, we performed *in situ* hybridization (ISH) for *Ccn1* on coronal forebrain sections at embryonic (E) 12.5, E14.5, E16.5, and E18.5. *Ccn1* was consistently detected in the germinal zone lining the lateral ventricle (LV), and the expression level was increased from E12.5 to E18.5 in the cortex (Figure 1A). Referring to the database from the Gene Expression Omnibus (GEO, Accession: GSE65000) (Florio et al., 2015), which offers gene expression profiling of subtypes of neural progenitor cells (NPCs), we found that *Ccn1* is highly enriched in apical RGCs (aRGCs) of both E14.5 mouse and 13wpc (weeks postconception) human brains, but expressed at minimal levels in basal RGCs (bRGCs), IPCs, and neurons (Figure S1A). The ISH result was further supported by an expression gradient map from the t-SNE clustering of E12∼E15 single-cell RNA-seq data of the cerebral cortex (Figure S1B) (Telley et al., 2019). Consistently, we detected CCN1 protein in PAX6+ (a marker for RGCs) cells and KI67+ (a marker for proliferative cells) cells by co-immunostaining for CCN1 and PAX6 or KI67 (Figure 1B).

**Figure 1.**
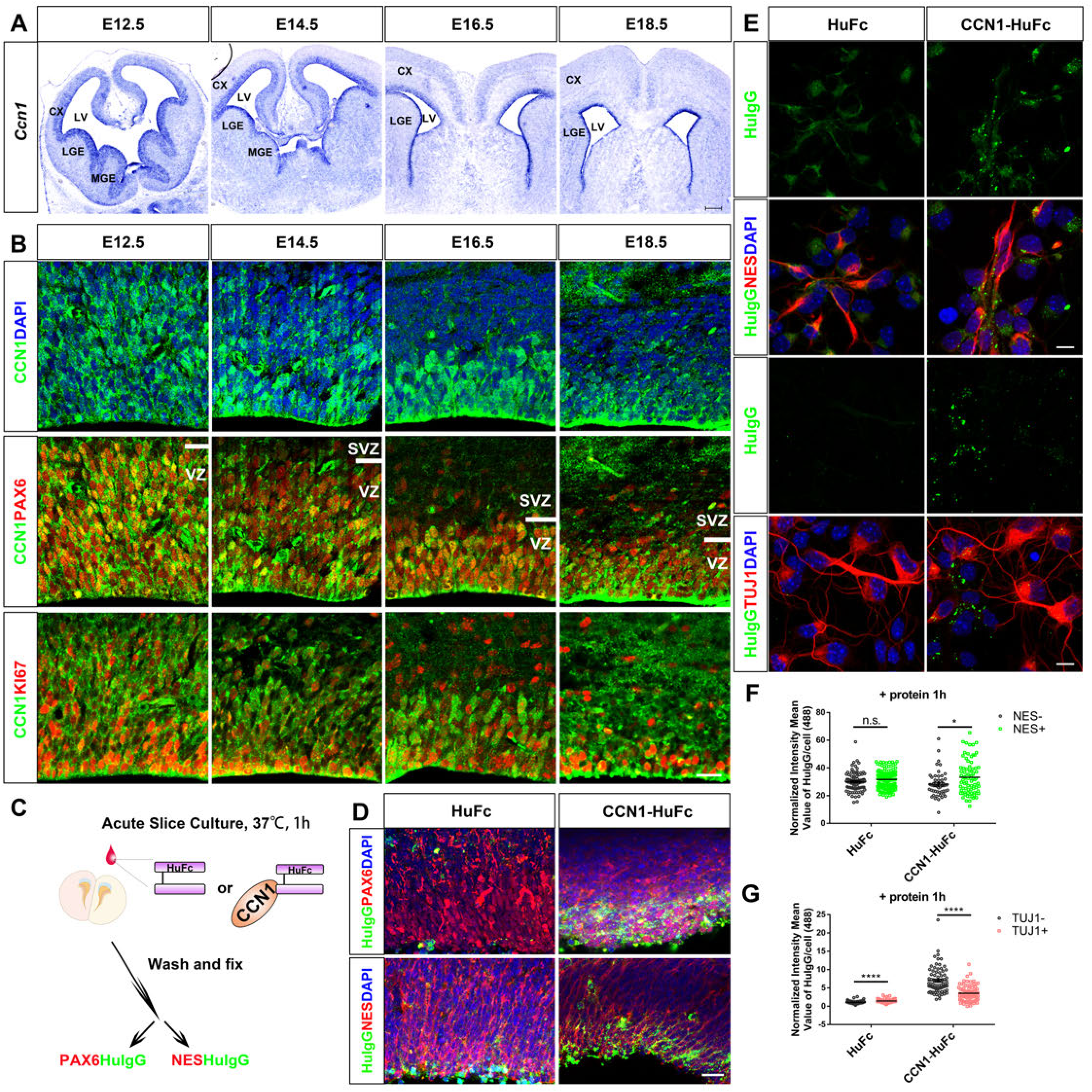
*Ccn1* is expressed in apical radial glial cells during development, and the majority of the matricellular CCN1 binds to radial glial cells. (A) *In situ* hybridization of *Ccn1* during embryonic developmental stages of E12.5, E14.5, E16.5, and E18.5. (B) Representative images of immunostaining with CCN1 (green) and PAX6 (second row, red) or KI67 (third row, red) at the indicated stages of the mouse cortices. (C) Schematic diagram of experimental design to clarify CCN1-HuFc binding cell types *ex vivo* (D) Representative images of IF staining of Human IgG (HuIgG, green) and PAX6 (upper, red) or NES (lower, red) on acute slice culture incubated with HuFc or CCN1-HuFc recombinant protein. (E) Representative images of co-staining of HuIgG (green) and NES (second row, red) or TUJ1 (fourth row, red), respectively, on primary NPCs dissociated from E13.5 cortices. The cells were stained after 1h incubation of HuFc or CCN1-HuFc, respectively, after culturing 24h. (F-G) Quantification of the normalized intensity mean value of HuIgG per cell (Alexa 488), which were added with HuFc or CCN1-HuFc proteins, respectively, for NES- and NES+ (F) and TUJ1- and TUJ1+ (G) cells from (E). The normalized intensity mean value (IMV) of each cell was calculated by subtracting the background IMV within the same view from the IMV of the cell measured. HuFc: N=87 for NES-, N=119 for NES+, P=0.068; CCN1-HuFc: N=50 for NES-, N=75 for NES+, P=0.0109. HuFc: N=73 for TUJ1-, N=66 for TUJ1+, P<0.0001; CCN1-HuFc: N=76 for TUJ1-, N=81 for TUJ1+, P<0.0001. DAPI (blue) was stained for the nucleus. LV: lateral ventricle, CX: cortex, LGE: lateral ganglionic eminence, MGE: medial ganglionic eminence. Scale bars: 200 μm for (A), 20 μm for (B and D), 10 μm for (E). Error bar columns represent mean ± SEM, unpaired student’s t test with Welch’s correction was used for the statistical test in (F and G), n.s. P>0.05, *P<0.05, **** P < 0.0001.

To test whether soluble CCN1 binds to NPCs in vivo, we added the commercialized HuFc (human IgG1 Fc peptide) or CCN1-HuFc recombinant protein into the E13.5 cortical slice culture. After wash and staining, we found that CCN1 protein remained mainly in the VZ region on the PAX6+ and NES+ (a marker for NSCs) cells, indicating CCN1 binds to RGCs *in vivo* (Figure 1C and 1D). Consistently, our analysis, through quantification of normalized intensity mean value of HuIgG, of CCN1 binding cell type assay *in vitro* further substantiated that the majority of the CCN1 proteins bind on the NES+ and TUJ1-RGCs rather than NES- and TUJ1+ neurons (Figure 1E-G, HuFc NES-: 29.90 ± 0.76, HuFc NES+: 31.69 ± 0.62, CCN1-HuFc NES-: 28.28 ± 1.23, CCN1-HuFc NES+: 33.25 ± 1.42; HuFc TUJ1-: 1.06 ± 0.05, HuFc TUJ1+: 1.43 ± 0.06, CCN1-HuFc TUJ1-: 7.01 ± 0.41, CCN1-HuFc TUJ1+: 3.55 ± 0.24). Furthermore, we constructed a plasmid expressing CCN1-EGFP fusion protein to visualize the localization of CCN1 protein. We introduced the plasmid into the cortical RGCs at E12.5 through *in utero* electroporation (IUE) and isolated the electroporated cells at E14.5 to culture for 2 days *in vitro* (Figure S1C). We found that CCN1-EGFP was distributed in a clustered punctate pattern along with the radial fiber of NES+ cells (Figure S1D).

### Conditional knockout of *Ccn1* accelerates RGC lineage progression and exerts premature cell cycle exit of NPCs, leading to reduced NSC pool size

As *Ccn1* knockout mice die of placental abnormalities and hemorrhage at early embryogenesis (Mo et al., 2002), to elucidate the function of CCN1 in embryonic RGCs *in vivo*, we generated CCN1 conditional knockout mouse lines (*Nestin*-Cre^+/-^/*Ccn1*^fl/fl^, CcKO) using the Cre-loxP system by crossing *Ccn1*^flox/flox^ mice with *Nestin*-Cre mouse lines. The specificity of Cre recombination has been verified in our previous study (Hu et al., 2017). Here, we confirmed the efficiency of the *Ccn1* deletion by ISH and RT-qPCR. The *Ccn1* mRNA was undetectable on E16.5 CcKO brain sections. However, the knockout efficiency at E13.5, especially E11.5, was insufficient (Figure S2A). The relative normalized expression of *Ccn1* in the cortices of the CcKO mice (0.47 ± 0.03) at E14.5 was significantly lower than that in the WT mice(1.00 ± 0.10). However, the difference between CcKO mice (0.80 ± 0.04) and the WT mice (1.00 ± 0.07) was mild in E11.5 (Figure S2B). These results suggested a late-onset complete knockout of *Ccn1* after the summit of neurogenesis. The survival or fertility of CcKO mice was normal compared to WT mice, and there was no apparent change in gross brain and body weight (data not shown).

To delineate the lineage progression of cortical NPCs, we co-stained PAX6 and EOMES (a.k.a. TBR2) on E15.5 brain sections. We calculated the percentage of PAX6+EOMES-RGCs, PAX6+EOMES+ transient IPCs, and PAX6-EOMES+ IPCs (Stenzel et al., 2014). The result showed the percentage of RGCs decreased (57.77% ± 0.02% for WT, 50.63% ± 0.79% for CcKO), and the transient IPCs increased (13.48% ± 0.90% for WT, 18.91% ± 1.24% for CcKO), while IPCs did not change (28.75% ± 0.92% for WT, 30.46% ± 1.89% for CcKO), which suggested that *Ccn1* deficiency leads to reduced number of RGCs and increased transition from RGCs to IPCs (Figure 2A and 2B). To determine whether the reduction in the number of RGCs was due to proliferation defects, we co-stained PAX6 and KI67 on E15.5 brain sections and analyzed the percentage of proliferative cells in VZ/SVZ subregions. We found that the percentage of proliferative cells in RGCs in the VZ/SVZ was significantly reduced (73.66% ± 1.18% for WT, 60.04% ± 2.07% for CcKO), which was mainly due to a decrease in the VZ (Figure 2C and 2D, 72.89% ± 1.21% for WT, 59.54% ± 3.05% for CcKO) rather than in the SVZ (79.64% ± 0.72% for WT, 65.60% ± 6.52% for CcKO), suggesting loss of *Ccn1* decreased the number of proliferating RGCs. We further performed abdominal Bromodeoxyuridine (BrdU) injection into E14.5 pregnant mice. We sacrificed the mice 24h later and co-stained BrdU with KI67 to determine the cell cycle exit index (BrdU+KI67-/BrdU+KI67+). There was an increase in the cell cycle exit index in CcKO (0.61 ± 0.04) compared to that in WT (0.46 ± 0.03) (Figure 2E-G). As a result, we found a decrease in the density of PAX6+ RGCs in VZ/SVZ by immunostaining on brain sections at E17.5 of CcKO compared to that of WT (Figure 2H and 2I, 182.1 ± 4.81 for WT, 161.7 ± 6.24 for CcKO). Hence, loss of *Ccn1* reduces RGC proliferation and increases cell cycle exit, consequently reducing RGCs.

**Figure 2.**
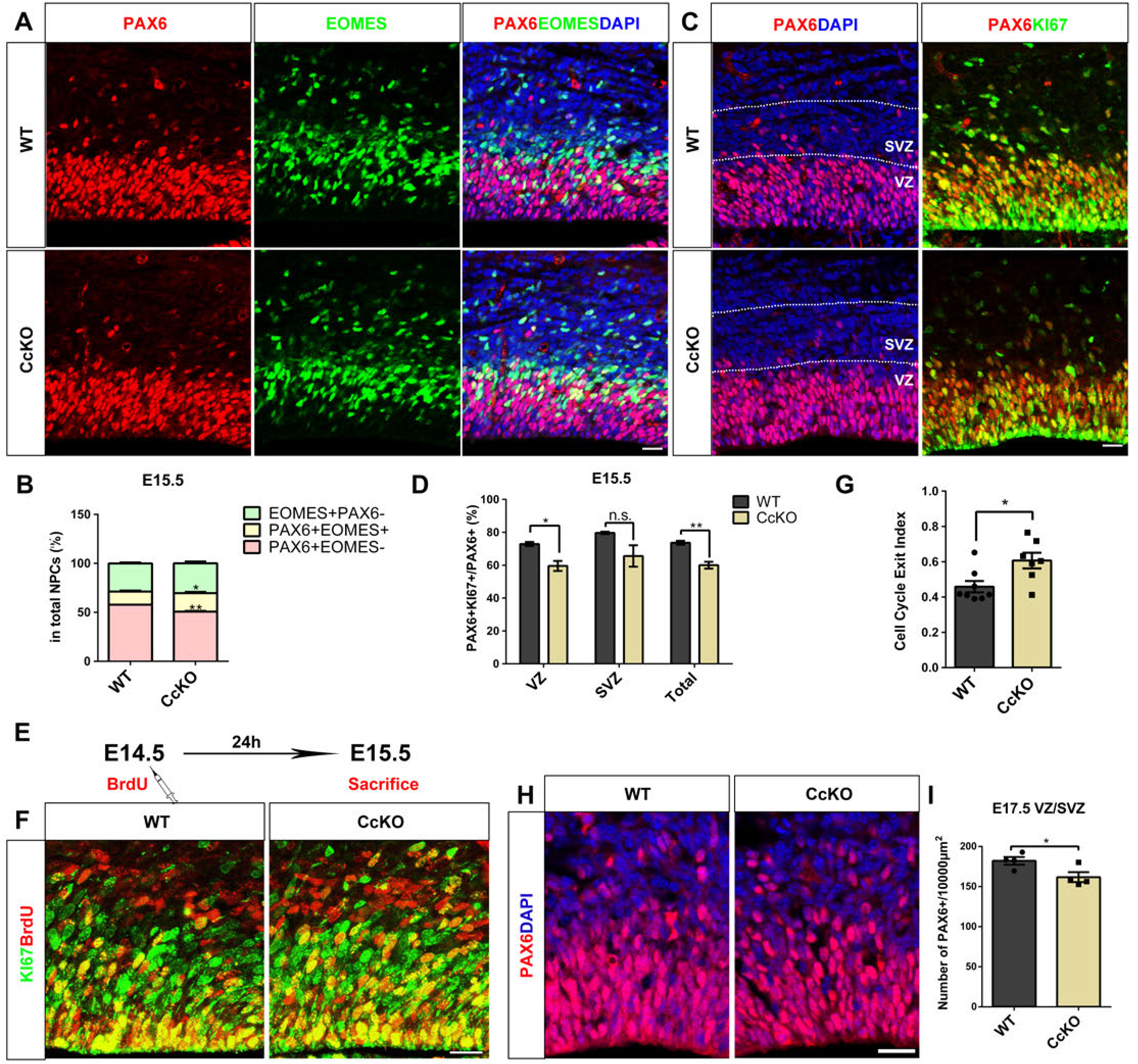
Conditional knockout of *Ccn1* accelerates lineage progression of RGCs and exerts premature cell cycle exit of NPCs, leading to reduction of NSC pool size. (A) Representative confocal images of E15.5 WT or CcKO mice brain coronal sections with PAX6 (red) and EOMES (green) double IF staining. (B) Statistical bar graphs for the percentages of PAX6+EOMES-, PAX6+EMOES+ and PAX6-EOMES+ subpopulation distribution of (A). N=3 for both WT and CcKO. P<0.01 (A, PAX6+EOMES-), P<0.05 (A, PAX6+EOMES+), P>0.05 (A, PAX6-EOMES+). (C) Images of E15.5 WT or CcKO mice brain coronal sections with PAX6 (red) and KI67 (green) double IF staining. Dashed lines depict the borders of VZ and SVZ. (D) Statistical bar graphs for the percentages of PAX6+KI67+ proliferative progenitors in PAX6+ progenitors in VZ, SVZ, or Total (VZ/SVZ) from (C). N=3 for both WT and CcKO. P=0.0346 (B, in VZ), P=0.1627 (B, in SVZ), P=0.009 (B, in total). (E-G) Representative images of BrdU (red) and KI67 (green) immunostaining (F) after the experimental procedure of BrdU injection at E14.5 and mice sacrifice 24h later (E), (G) shows the statistical bar graph for cell cycle exit index (BrdU+KI67-/BrdU+KI67+) of WT or CcKO mice cortical NPCs. WT: N=8, CcKO: N=7, P=0.0203. (H) Representative images of PAX6 (green) immunostaining on the coronal forebrain sections of WT and CcKO mice at E17.5. (I) Quantification of the number of PAX6+ RGCs within 10000 μm^2^ area in WT and CcKO mice, WT: N=4, CcKO: N=4, P=0.0439. DAPI (blue) was stained for the nucleus. VZ: ventricular zone, SVZ: subventricular zone. Scale bars: 20 μm for (A, B, and F), 20 μm for (H). Error bar columns represent mean ± SEM, two-way ANOVA was used for the statistical test in (C), unpaired student’s t test with Welch’s correction was used for the statistical test in (D, G, and I), *P<0.05; ** P < 0.01.

After figuring out the disturbance of NPCs caused by *Ccn1* deletion, we hoped to determine whether the projection neurons sequentially produced from NPCs during development are perturbed in the result. Thus, we stained SATB2 (a Layer II-VI projection neuron marker) on brain sections at postnatal day 30 (P30) (Figure S2C). The density of SATB2+ neurons in Layer II-VI was not significantly changed in CcKO compared to WT (391.7 ± 14.54 /10 mm^2^ for WT, 337.0 ± 23.63 /10 mm^2^ for CcKO) was significantly decreased (Figure S2D). Also, there was no significant difference in the thickness of cortex (1.17 ± 0.02 mm for WT, 1.16 ± 0.06 mm for CcKO) between WT and CcKO mice (Figure S2E).

### Loss of *Ccn1* disturbs NSC maintenance, resulting in premature neuronal differentiation at the expense of perinatal gliogenesis

To further dissect the effect of *Ccn1* deletion on radial glial lineage, we utilized the *PiggyBac* transposon system, which induces nonviral genomic integration of transgenes in transfected cells, to reveal the complete lineage of cortical RGCs and their progenies, including both neurons and glial cells (Chen et al., 2014). We electroporated the *PiggyBac*-pCAG-IRES-EGFP (we designated as PB-Ctrl) plasmid into the WT or CcKO cortex at E14.5. Then we injected BrdU into the abdominal cavity of the pregnant mice after 24h and analyzed the distribution and the cell types of GFP+ cells in the cortices at P10 (Figure 3A). We calculated the normalized relative GFP intensity mean in VZ/SVZ, corpus callosum (CC, glial cells in major), deep cortical plate (DCP), and upper cortical plate (UCP), respectively. Remarkably, the GFP+ cells in the VZ/SVZ and CC of the *Ccn1*-deficient neocortex was completely lost compared to WT (Figure 3B’ and 3C, VZ/SVZ: 18.20% ± 2.70% for WT, 4.44% ± 1.63% for CcKO; CC: 13.28% ± 0.76% for WT, 1.87% ± 0.88% for CcKO; DCP: 12.81% ± 0.60% for WT, 5.29% ± 2.25% for CcKO; UCP: 55.71% ± 2.86% for WT, 88.40% ± 4.31% for CcKO), and the percentage of the GFP+SATB2+ neurons were dramatically increased in CcKO UCP compared to WT (Figure 3B’’, 3B’’’, and 3D, 72.67% ± 1.81% for WT, 88.77% ± 0.45% for CcKO), which was confirmed by the increase of GFP+CUX1+ neurons of CcKO UCP in contrast to that of the WT (Figure S3A and S3B, 68.41% ± 1.84% for WT, 81.99% ± 2.51% for CcKO). Notably, the decreased GFP+SATB2-cells with dense bushy short processes were morphologically like astrocytes (Siddiqi et al., 2014), whose morphology was largely different from GFP+SATB2+ pyramidal neurons (Figure 3B’’’). We also found that although the percentage of GFP+SATB2+BrdU+ cells generated at E15.5 in all GFP+SATB2+ neurons was similar in the cortex of CcKO to that in control (Figure S3C and S3D, 19.00% ± 3.55% for WT, 22.71% ± 2.66% for CcKO), the location of GFP+SATB2+ neurons in CcKO was mostly restricted to a narrow band below layer II/III: the distribution of GFP+SATB2+ neurons in the lower part of the UCP was increased, but diminished in the upper part of UCP of CcKO cortex (Figure 3B’’ and 3E, in WT: Layer I: 0.07% ± 0.07%, Layer II/III: 62.14% ± 1.58%, Layer IV: 37.80% ± 1.52%; in CcKO: Layer I: 0, Layer II/III: 33.66% ± 4.67%, Layer IV: 66.34% ± 4.67%), suggesting that E14.5 RGCs in CcKO gave rise to SATB2+ neurons within a shorter time period. These results indicate that *Ccn1* is required for maintaining the cortical NSC pool and loss of *Ccn1* leads to premature neuronal differentiation and depletion of glial progenitor cells.

**Figure 3.**
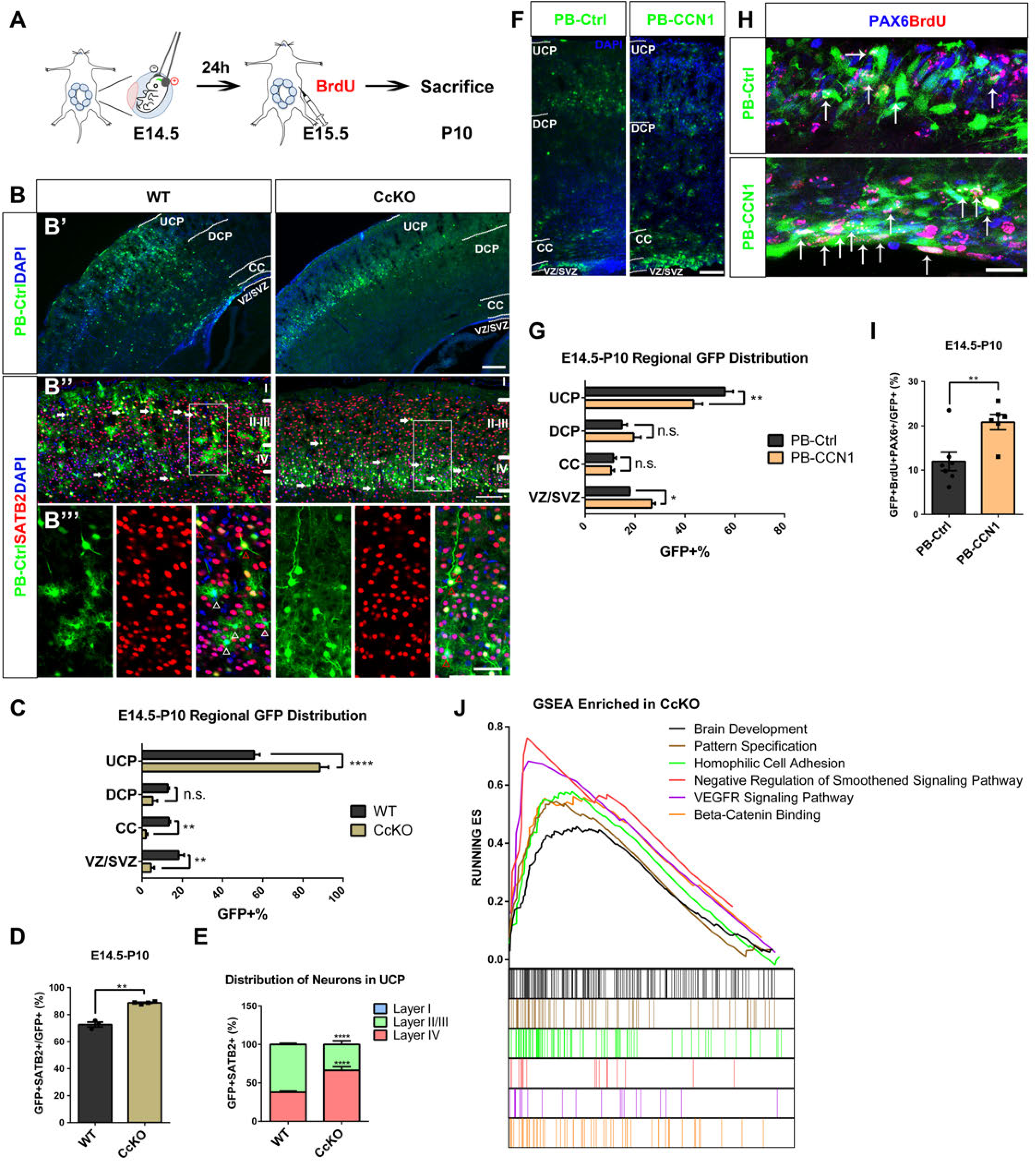
Loss of *Ccn1* causes NSC depletion and interferes with brain developmental genes and multiple signaling pathways, resulting in premature neuronal differentiation at the expense of perinatal gliogenesis. (A) Schematic diagram of the experimental design of IUE (E14.5 → P10) with plasmids in pregnant mice and the following BrdU injection (E15.5) for Figure 3 (B, G, and I) and Figure S5. (B) Representative images of the distribution of the cortical GFP+ (PB-Ctrl) cells after IUE for WT or CcKO mice, white dashed lines indicate the borders of UCP, DCP, CC, and VZ/SVZ (B’). The zoom-in images of the cortical plate with SATB2 (B’’, red) immunostaining on the brain sections from (B’), arrows point to GFP+SATB2+ cells (A). Zoom-in images of the white box in (B’’), white arrowheads point to the GFP+SATB2-glial cells with highly ramified morphology, red arrowheads point to GFP+SATB2+ pyramidal neurons (B’’’). (C-E) Statistical bar graphs for (B). The percentage of GFP+ cells labeled by PB-Ctrl in WT or CcKO mice cortices within UCP, DCP, CC, and VZ/SVZ, respectively (C). The percentage of SATB2+ neurons in UCP GFP+ cells in the WT or CcKO mice cortex (D). The percentage of GFP+SATB2+ cells’ distribution in the layers of the UCP (E). WT: N=3, CcKO: N=4, P<0.01 (C, VZ/SVZ), P<0.01 (C, CC), P<0.0001 (C, UCP), P= 0.0092 (D), P<0.0001 (E, Layer II/III and Layer IV). (F) Representative images of the distribution of GFP+ cells from the cortices electroporated with PB-Ctrl or PB-CCN1 plasmids through procedures in (A), white dashed lines indicate the borders of UCP, DCP, CC, and VZ/SVZ. (G) Statistical bar graph for the distribution percentage of GFP+ cells within the UCP, DCP, CC, and VZ/SVZ subregions, respectively, of the cortices electroporated with the indicated plasmids. PB-Ctrl: N=4, PB-CCN1: N=4, P<0.05 (VZ/SVZ), P<0.01 (UCP). (H) Zoom-in images of PB-Ctrl or PB-CCN1 (green) brain sections immunostained with PAX6 (blue) and BrdU (red), from SVZ of (F). Arrows point to PAX6+BrdU+GFP+ cells. (I) Quantification for the percentage of PAX6+BrdU+GFP+ label-retaining progenitor cells from E15.5 in GFP+ cells within VZ/SVZ of (F). PB-Ctrl: N=7, PB-CCN1: N=6, P=0.0077. (J) GSEA enrichment score line chart of genes ranked by Fold Change value, the bar codes at the bottom are the ranked genes belonging to the gene sets in corresponding colors. X-axis: rank in the gene list, Y-axis: running enrichment score (ES). DAPI (blue) was stained for the nucleus. UCP: upper cortical plate (Layer I-IV); DCP: deep cortical plate (Layer V-VI); CC: corpus callosum. Scale bars: 200 μm for images in (B’), 100 μm for images in (B’’) and (F), 50 μm for (B’’’), 20 μm for (H). Error bar columns represent mean ± SEM, two-way ANOVA was used for statistical test in (C, E, and G), unpaired student’s t test with Welche’s correction was used for statistical test in (D and I), *P<0.05; ** P < 0.01; *** P < 0.001; **** P < 0.0001.

In addition to loss-of-function analysis, we explored the gain-of-function phenotype of CCN1 using PB-Ctrl and *PiggyBac*-pCAG-CCN1-IRES-EGFP (we designated as PB-CCN1) plasmids following the previous procedures (Figure 3A). Overexpression of PB-CCN1 was efficient, shown by western blot using an anti-CCN1 antibody (Figure S3E). We performed a statistical analysis of the percentage of the GFP+ cells of PB-CCN1 and PB-Ctrl and found more GFP+ cells distributed in VZ/SVZ after CCN1 overexpression compared to control (Figure 3F and 3G, VZ/SVZ: 17.92% ± 0.10% for WT, 26.69% ± 1.46% for CcKO; CC: 11.28% ± 1.20% for WT, 10.36% ± 1.30% for CcKO; DCP: 14.75% ± 2.09% for WT, 19.44% ± 2.90% for CcKO; UCP: 56.05% ± 3.26% for WT, 43.51% ± 3.61% for CcKO). Then we co-stained BrdU and PAX6 on indicated brain sections and compared the percentage of PAX6+BrdU+GFP+ label-retaining progenitor cells since E15.5 in GFP+ cells within SVZ in PB-CCN1 (20.83% ± 1.72%) and PB-Ctrl (11.98% ± 2.10%) (Figure 3H and 3I). To sum up, the BrdU-incorporated NPCs retained in the SVZ were increased if CCN1 was constitutively overexpressed, which further substantiated that CCN1 maintains NSCs during cortical development.

### CCN1 regulates brain developmental genes and multiple signaling pathways

To determine the underlying molecular processes and biological functions in NSC lineage impacted by CCN1, we performed whole-genome RNA-sequencing (RNA-seq) analysis of NSC lineage cells labeled by IUE GFP-expressing constructs in CcKO and control cortex. Three days after IUE, by fluorescence-activated cell sorting (FACS), we collected GFP+ cells, including the initial electroporated cells and their progeny. We determined the differentially expressed genes (DEGs) between GFP+ cells from CcKO and control (Figure S3F). The significant DEGs were defined as genes with the absolute value of Fold Change>1.2 and P-value<0.05. There were 1712 significantly UP-regulated genes and 1357 significantly DOWN-regulated genes, *Ccn1* was downregulated significantly (Figure S3G). We performed the gene set enrichment analysis (GSEA) to clarify the enriched biological functions of transcriptomic differences combining the ranked fold change of gene expression from the functional gene sets (Mootha et al., 2003; Subramanian et al., 2005). In addition to the enrichment of genes related to brain development and pattern specification in CcKO mice, the negative regulation of the Smoothened signaling pathway and the VEGFR signaling pathway were the two pathways with the highest enrichment scores. The gene set of beta-catenin binding was also significantly enriched in CcKO mice (Figure 3J). Beta-*Catenin* has been reported to be crossroads of multiple signaling pathways of niche factors, such as Notch, Wnt (Kim et al., 2009; Mao et al., 2009; Shimizu et al., 2008; Zhang et al., 2010). In brief, *Ccn1* deficiency perturbed the activities of multiple niche factor signaling pathways that regulate NSC maintenance.

### CCN1 interacts with GPC4 core protein on the membrane of radial glial cells

As a secreted matricellular protein, CCN1 is capable of binding to multiple membrane proteins and extracellular matrix proteins (Lau, 2011). To pinpoint the binding partners of CCN1 in the developing forebrain, we designed a pull-down assay using recombinant CCN1-HuFc or HuFc control protein to enrich proteins that bind to CCN1 in the E13.5 forebrain protein lysate (Figure S4A). The silver staining of two repeats of pull-down proteins in the HuFc or CCN1-HuFc group showed distinctive differential protein bands between the two groups (Figure S4B). By mass-spectrometry analysis, we identified 258 proteins that were specifically detected in the CCN1-HuFc pull-down group after filtering out nuclear-localized and ribosome-binding proteins (Figure 4A). To focus on the processes involved in extracellular-to-intracellular signaling, based on the enrichment intensity of interacting proteins and gene ontology (GO) analysis, we selected proteins in “receptor-mediated endocytosis” and “heparan sulfate-glycosaminoglycan (HS-GAG) biosynthesis” for further verification (Figure 4B). Particularly, we tested the interaction between CCN1 and members of the heparan sulfate proteoglycan (HSPG) subfamily (glypicans) through immunoprecipitation (IP). We found that CCN1 was evidently bound to Glypican-4 (GPC4) while its binding to other glypicans (GPC1,2,6) were barely detectable (Figure 4C). Consistently, CCN1 and GPC4 were detected in close proximity in Neuro2a (N2a) cells transfected with both CCN1-RFP and GPC4-EGFP fusion protein constructs. As expected, we found that GPC4-EGFP was localized around the cell membrane where dots of extracellular CCN1-RFP were (Figure S4C).

**Figure 4.**
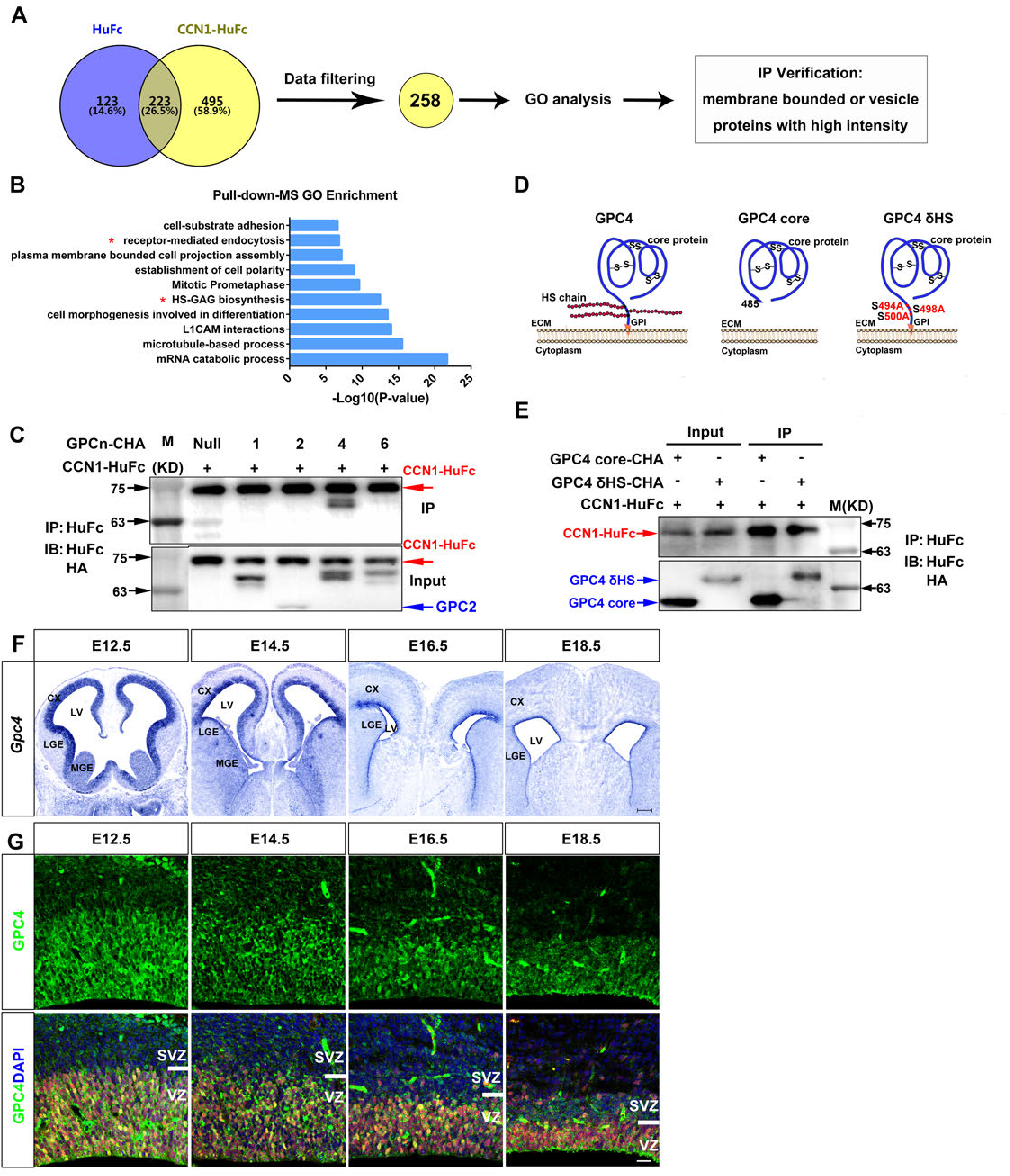
CCN1 interacts with GPC4 on the membrane of radial glial cells. (A) Schematic diagram of data processing design for pull-down-MS data, including the Venn diagram of indicated bait proteins and the data filtering strategies for IP verification. (B) Bar chart for GO enrichment of CCN1 interacting proteome in the embryonic forebrain. Red asterisks indicate the enriched GO term to which our selected IP verification membrane proteins belong. (C) Western blot of input (lower) or IP (upper) samples for CCN1-HuFc with GPC1/2/4/6-CHA, respectively. (D) Diagrams for GPC4 core protein and GPC4 δHS mutant construction. (E) Western blot of input (lane 1 and 2) or IP (lane 3 and 4) samples for CCN1-HuFc with GPC core-CHA or GPC4 δHS-CHA, respectively. (F) *In situ* hybridization of *Gpc4* during indicated embryonic developmental stages. (G) Images of IF staining with GPC4 (green) and PAX6 (second row, red) at the indicated stages of the mouse cortex. Scale bars: 200 μm for (F), 20 μm for (G).

The functional domains of GPC family proteins consist of a compact globular core protein domain, the C-terminal glycosylphosphatidylinositol (GPI) cell membrane anchor, and the X-G-X-G-S-X motif of serine covalent binding sites of heparan sulfate (HS) chains adjacent to C-terminal (Fico and Dono, 2012). We further constructed the GPC4 core-CHA protein (amino acid, hereafter, aa.1-485) and GPC4 δHS-CHA (S494A, S498A, S500A) mutant protein and tested the interactions between these mutants and CCN1-HuFc through IP. It turned out that GPC4 interacts with CCN1 through its core protein domain independent of HS chains or GPI anchor (Figure 4D and 4E). Intriguingly, GPCs, especially GPC4, are clustered separable on the cell membrane according to two types of HS chain modifications, namely N-acetal-rich or N-sulfo-rich, and the clustering is generalized across cell types (Mii et al., 2017), reminding us of the punctate dots of CCN1 we observed (Figure S1D). Thus, we co-stained HuIgG and N-sulfo HS (HepSS-1) on RGCs incubated 0.5h with CCN1-HuFc after 24h culturing from E13.5 dissociated cortical cells. We found that the localization of the majority of the CCN1 puncta is separated from that of the HepSS-1 puncta (Figure S4D and S4E, 35.33% ± 5.18% for CCN1+HepSS-1+ puncta, 64.67% ± 5.18% for CCN1+HepSS-1-puncta), indicating the internalized CCN1 in the cytoplasm mediated by N-sulfo-rich HS chain GPC4 clusters and the remaining CCN1 mediated by N-acetal-rich HS chain GPC4 clusters on the cell membrane, which was consistent with the characterization of those two types of GPC4 clusters (Mii et al., 2017).

Previous studies have shown that *Gpc4* is expressed in the nervous system during development (Hagihara et al., 2000; Ybot-Gonzalez et al., 2005). Among all six *Gpcs*, *Gpc4* showed the most similar expression pattern to *Ccn1* on the cortex (Figure S4F) (Telley et al., 2019). To further confirm the expression pattern of GPC4 in the embryonic telencephalon, we performed ISH with the *Gpc4* RNA probe on brain sections from E12.5 to E18.5 mice. Similar to *Ccn1*, *Gpc4* was mainly expressed in the germinal zone lining the lateral ventricle throughout development. However, different from the ventral-high dorsal-low expression pattern of CCN1, we found *Gpc4* was prominently expressed in the developing cerebral cortex and the lateral ganglionic eminence (LGE) but much lower in the medial ganglionic eminence (MGE) (Figure 4F). Co-immunostaining of GPC4 and PAX6 exhibited that GPC4 was mainly localized to the cell membrane of PAX6+ RGCs during cortical development (Figure 4G). These results suggest that interaction between CCN1 and GPC4 is likely involved in niche signaling during RGC development.

### CCN1 is required for GPC4 to maintain NSCs, which may be through regulating the Shh signaling pathway and cell cycle related-pathways

To understand the function of GPC4 during cortical development, we electroporated the *PiggyBac*-pCAG-GPC4-IRES-EGFP (designated as PB-GPC4), PB-CCN1, and PB-Ctrl plasmids, respectively, at E14.5. Analyses of E17.5 cortices after IUE showed that overexpression of *Gpc4* and *Ccn1* similarly increased the percentage of SOX2+ NPCs in GFP+ cells localized in VZ/SVZ (41.19% ± 2.58% for PB-Ctrl, 51.45% ± 0.18% for PB-CCN1, 50.21% ± 0.41% for PB-GPC4) (Figure 5A and 5B). Interestingly, both CCN1 and GPC4 significantly reduced the percentage of proliferating SOX2+ cells (SOX2+GFP+KI67+) in VZ/SVZ SOX2+GFP+ cells (53.10% ± 3.25% for PB-Ctrl, 43.63% ± 4.28% for PB-CCN1, 46.68% ± 2.14% for PB-GPC4) (Figure 5A and 5C). These results suggested that GPC4, similar to CCN1, promotes the maintenance of NSCs in a non-proliferative state. Meanwhile, GPC4 peculiarly increases the ratio of SOX2+ NPCs in the SVZ to that in the VZ, in contrast to the comparable ratio of CCN1 compared to the control (0.25 ± 0.03 for PB-Ctrl, 0.24 ± 0.06 for PB-CCN1, 0.41 ± 0.05 for PB-GPC4) (Figure 5A and 5D).

**Figure 5.**
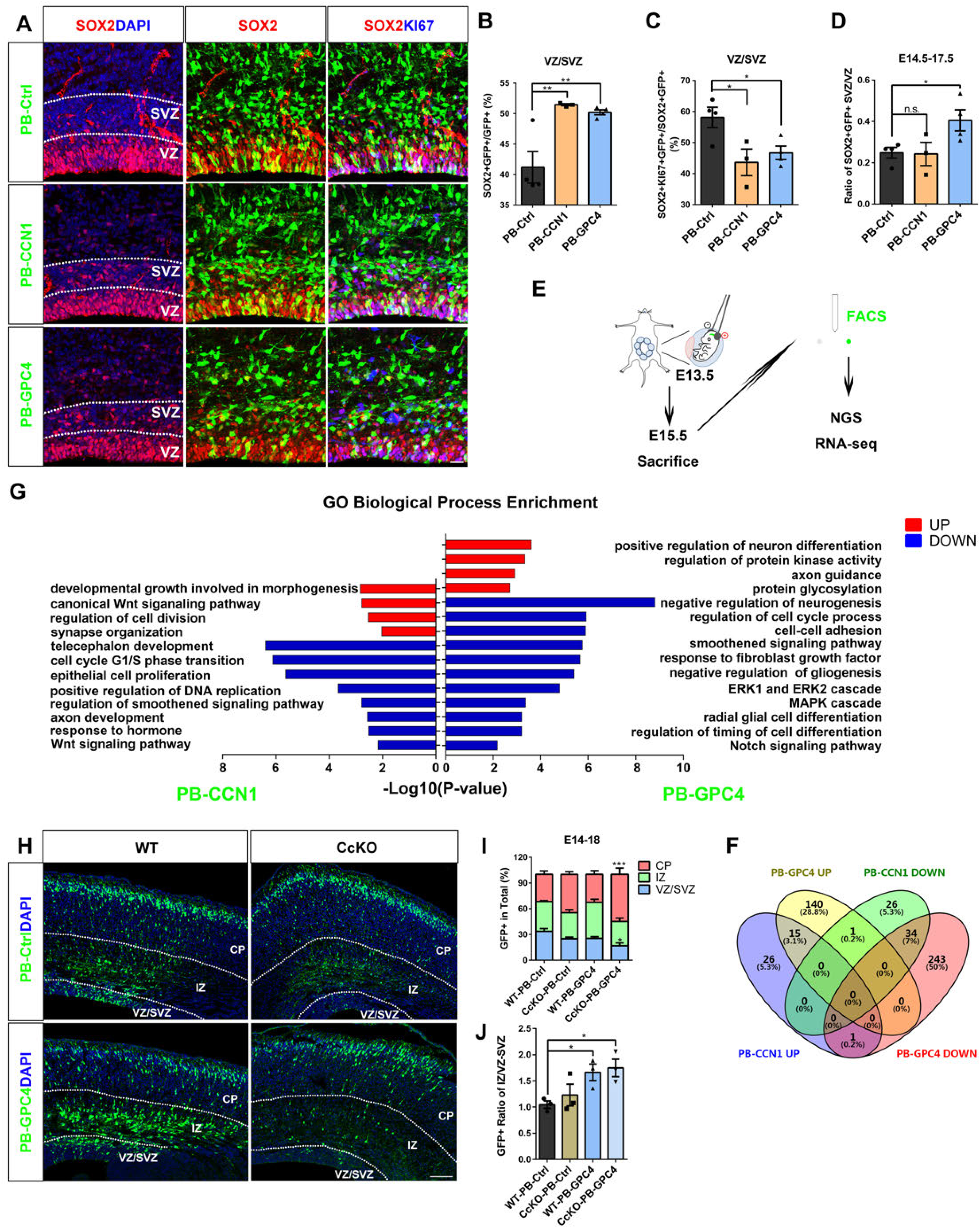
CCN1 is required for GPC4 to maintain NSCs and maybe through regulating the Shh signaling pathway and cell cycle-related pathways, while GPC4 induces more progenitors in SVZ distinct from CCN1. (A) Representative images of SOX2 (red) and KI67 (third column, blue) co-stained at E17.5 after IUE of PB-Ctrl, PB-CCN1, or PB-GPC4 (green) at E14.5. White dashed lines indicate the border of VZ and SVZ. (B-D) Statistical bar graphs for (A). The percentage of SOX2+GFP+ NPCs in GFP+ cells in VZ/SVZ for PB-Ctrl, PB-CCN1, or PB-GPC4 experimental groups (B). The percentage of KI67+SOX2+GFP+ proliferative NPCs in SOX2+GFP+ cells for corresponding groups (C). The ratio of SOX2+GFP+ NPCs in SVZ to VZ in respective groups (D). PB-Ctrl: N=4, PB-CCN1: N=3, PB-GPC4: N=4. P=0.0046 (B), P=0.0269 (C), P= 0.9294 (D, PB-CCN1 *vs.* PB-Ctrl), P=0.0471 (D, PB-GPC4 *vs.* PB-Ctrl). (F) Schematic diagram of experimental design of IUE (E13.5 ◊ E15.5) with PB-Ctrl, PB- CCN1, or PB-GPC4 plasmids into the cortex, purification of GFP+ cells by FACS, and the following RNA-seq analysis. (G) Venn diagram for UP or DOWN-regulated DEGs of PB-CCN1 or PB-GPC4. Compared to PB-Ctrl, we acquired 42 UP-regulated and 61 DOWN-regulated DEGs in the PB-CCN1 group and 156 UP-regulated and 278 DOWN-regulated DEGs in the PB-GPC4 group in parallel. There were 15 overlap genes up-regulated concordantly and 34 genes down-regulated concordantly by PB-CCN1 and PB-GPC4. (H) Bar chart for GO biological process enrichment of UP or DOWN-regulated DEGs of PB-CCN1 or PB-GPC4 comparing to PB-Ctrl, respectively. (I) Representative images of the E18.5 cortex after IUE with PB-Ctrl or PB-GPC4 (green) at E14.5 in WT or CcKO embryos. Dashed lines indicate VZ/SVZ, IZ, and CP border. (I and J) Statistical bar graph for (H). The distribution percentage of GFP+ cells in indicated subregions of the WT or CcKO cortices (I). The ratio of GFP+ cells’ distributed in IZ to VZ/SVZ (J). WT-PB-Ctrl: N=3, CcKO-PB-Ctrl: N=3, WT-PB-GPC4: N=3, CcKO-PB-GPC4: N=3. P<0.05 (I, VZ/SVZ of CcKO-PB-GPC4 vs. WT-PB-Ctrl), P<0.001(I, CP of CcKO-PB-GPC4 vs. WT-PB-Ctrl); P<0.05 (J, CcKO-PB-GPC4 vs. WT-PB-Ctrl, one-way ANOVA), P=0.043 (J, WT-PB-GPC4 vs. WT-PB-Ctrl, unpaired student’s t test). DAPI (blue) was stained for the nucleus. Scale bars: 20 μm for (A), 100 μm for (H). Error bar columns represent mean ± SEM, one-way ANOVA was used for the statistical test in (B, C, and J), unpaired student’s t test with welch’s correction was used for the statistical test in (D and J), two-way ANOVA was used for the statistical test in (I), *P<0.05; ** P < 0.01; *** P < 0.001; n.s. P>0.05.

To test the hypothesis that CCN1 and GPC4 are involved in common signaling pathways, we compared the molecular targets affected by CCN1 and GPC4. We analyzed the RNA-seq data from FACS purified cells dissociated from E15.5 cortices after IUE of PB-CCN1, PB-GPC4, or PB-Ctrl plasmid, respectively, at E13.5 (Figure 5E). Compared to PB-Ctrl, PB-CCN1 and PB-GPC4 efficiently increased the FPKM level of *Ccn1* and *Gpc4*, respectively. Interestingly, while *Ccn1* did not change the expression level of *Gpc4*, overexpression of *Gpc4* down-regulated the expression of *Ccn1* (Figure S5A). Here, we defined the significant DEGs as genes with the absolute value of Fold Change>1.2 and P-value<0.05. Compared to PB-Ctrl, the majority of the overlapping DEGs were concordantly regulated by PB-CCN1 and PB-GPC4 rather than oppositely (Figure 5F). GO enrichment analysis of DEGs revealed that both PB-CCN1 and PB-GPC4 significantly down-regulated cell cycle-related pathways, such as the regulation of the cell cycle process and cell cycle G1/S transition (Figure S5B), consistent with the cellular phenotypes shown in Figure 5A-C. We identified the Smoothened signaling pathway as the common pathway affected by both CCN1 and GPC4. We also noticed CCN1 affected Wnt signaling pathway-related GO terms, while PB-GPC4 down-regulated GO terms related to Notch and FGF signaling pathways (Figure 5G), suggesting the interaction of GPC4 and CCN1 could confer cross-linking multiple signaling pathways. Indeed, genes involved in Shh, Wnt, Notch, or FGF signaling pathways, such as *Gas1*, *Sall3*, and *Notch2*, were consistently regulated by CCN1 and GPC4 (Figure S5C). GAS1 protein has been reported to be a co-receptor of SHH. It was induced by dorsal Wnt to antagonize the activity of ventral SHH during the early dorsal-ventral formation of somite (Lee et al., 2001). Intriguingly, we found *Gas1* is expressed in the germinal zone of the forebrain with dorsal-high to ventral-low gradient similar to *Gpc4* (Figure S5D, from *Allen Brain Atlas*).

To assess the functional associations between CCN1 and GPC4, we analyzed the lineage progression of RGCs in WT or CcKO brain by IUE with PB-Ctrl or PB-GPC4 plasmids at E14.5 and examined brain sections at E18.5. We quantified the percentage of GFP+ cells in the VZ/SVZ, IZ, and CP to indicate the NSC maintenance in VZ/SVZ and neuronal differentiation in postmitotic IZ and CP (Diaz-Alonso et al., 2017). We found that overexpression of GPC4 significantly affected the distribution of GFP+ cells in the absence of CCN1 (CcKO-PB-GPC4) compared to WT-PB-Ctrl, the percentage of differentiated neurons was increased in either CcKO-PB-Ctrl (significant when compared in Figure 6D) or CcKO-PB-GPC4 cortices (Figure 5H and 5I, WT-PB-Ctrl: CP: 31.61% ± 4.02%; CcKO-PB-Ctrl: CP: 44.63% ± 3.14%; WT-PB-GPC4: CP: 32.51% ± 4.27%; CcKO-PB-GPC4: CP: 54.87% ± 7.30%). Meanwhile, the ratio of GFP+ cells in the IZ to VZ/SVZ either in WT-PB-GPC4 or CcKO-PB-GPC4 cortices was increased (WT-PB-Ctrl: 1.05 ± 0.07, CcKO-PB-Ctrl: 1.23 ± 0.21, WT-PB-GPC4: 1.66 ± 0.16, CcKO-PB-GPC4: 1.75 ± 0.17) (Figure 5J). Further immunohistological analysis of GFP+SOX2+ NPCs in VZ or SVZ showed the ratio of GFP+SOX2+ cells in SVZ to VZ either in WT-PB-GPC4 or CcKO-PB-GPC4 cortices was increased (Figure S5E and S5F, WT-PB-Ctrl: 0.32 ± 0.05, CcKO-PB-Ctrl: 0.40 ± 0.06, WT-PB-GPC4: 0.59 ± 0.07, CcKO-PB-GPC4: 0.57 ± 0.06), which was consistent with Figure 5A and 5D. Together, these results suggest that GPC4 facilitates the dislocated NPCs in the SVZ and GFP+ cells in IZ independent of CCN1, while CCN1 is required for GPC4 to maintain NSCs.

**Figure 6.**
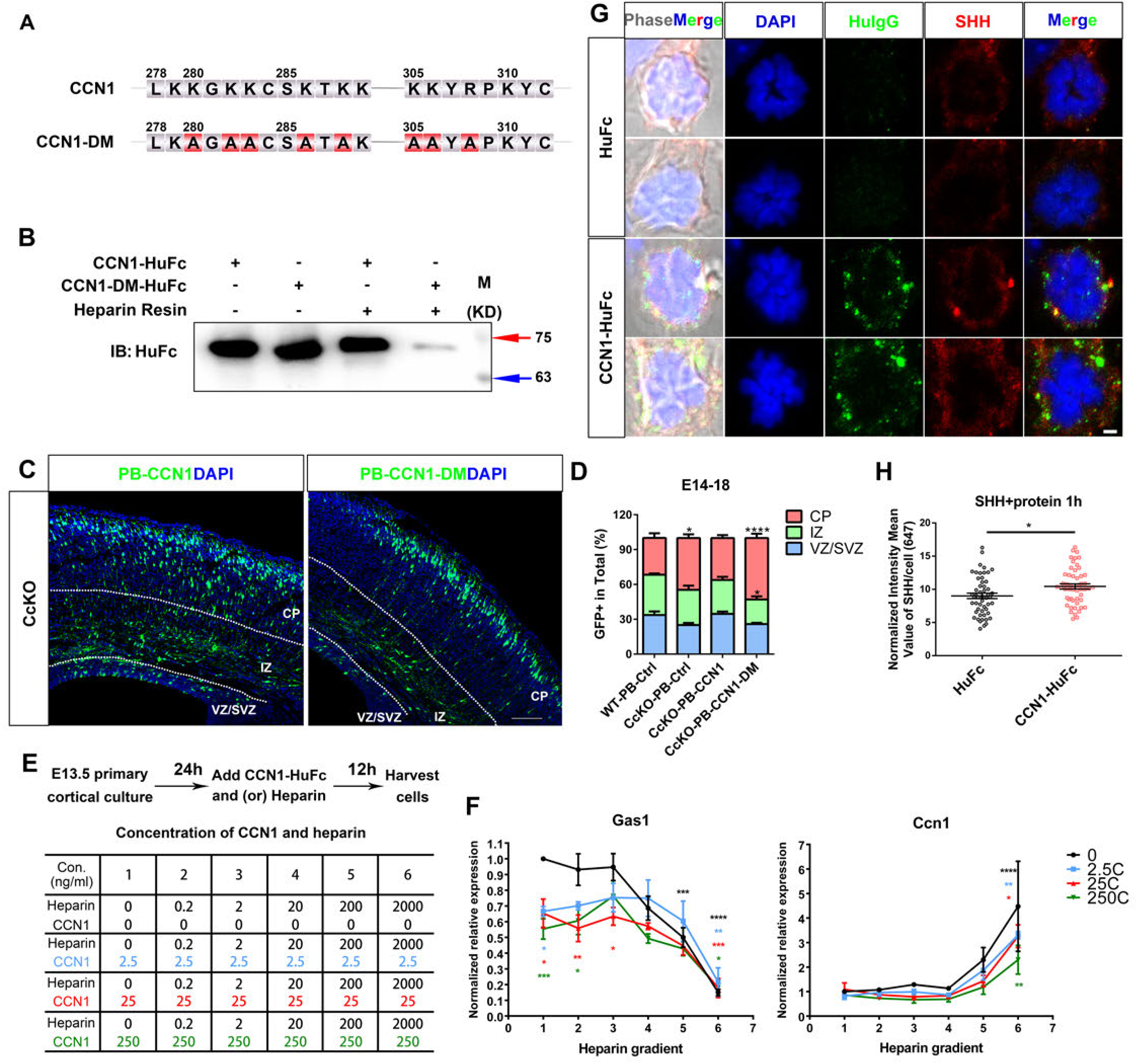
Heparin-binding is necessary for CCN1 to maintain neural stem cells, and they cooperate to regulate the Shh signaling pathway activities. (A) Mutation strategy for the heparin-binding mutant PB-CCN1-DM construction. (B) Western blot of enrichment of CCN1-HuFc or CCN1-DM-HuFc by heparin resin, lanes without heparin resin were inputs harvested 48h post-transfection in 293FT cell line. (C) Representative images of the E18.5 cortex after IUE of PB-CCN1 or PB-CCN1-DM (green) at E14.5 in CcKO embryos. Dashed lines indicate VZ/SVZ, IZ, and CP border. (D) Statistical bar graph for (C) and Figure 5 (H) PB-Ctrl groups. The percentage of GFP+ cells’ distribution in indicated sub-regions of the WT or CcKO cortex. WT-PB-Ctrl: N=3, CcKO-PB-Ctrl: N=3, CcKO-PB-CCN1: N=4, CcKO-PB-CCN1-DM: N=3. P<0.05 (CP of CcKO-PB-Ctrl *vs.* WT-PB-Ctrl), P<0.0001 (CP of CcKO-PB-CCN1-DM *vs.* WT-PB-Ctrl), P<0.05 (IZ of CcKO-PB-CCN1-DM *vs.* WT-PB-Ctrl). (E) Experimental design of the concentrations applied for the CCN1-HuFc recombinant protein and heparin to the E13.5 primary cortical cells cultured at div1 (day *in vitro*). The treated cells were harvested after 12h. (F) Statistical line charts of normalized relative *Gas1* and *Ccn1* endogenous expression by RT-qPCR. N=4. *Gapdh* was used as the reference gene, and each normalized gene expression was divided by no heparin and no CCN1-HuFc group within each independent repeat. The asterisks below the broken line indicate the significance of different CCN1-HuFc protein concentrations comparing the group without CCN1-HuFc protein at the same heparin concentration. The asterisks above the broken line indicate the significance of different heparin concentrations comparing the group without heparin at the same CCN1-HuFc protein concentration. The color of asterisks corresponds to the CCN1-HuFc protein concentration. (G) Representative images of the co-immunostaining of HuIgG (green) and SHH (red) on RGCs 24h after dissociated from E13.5 cortices and 1h incubation with SHH and HuFc or CCN1-HuFc, respectively. Both HuIgG and SHH were shown with the same parameters with Zeiss ZEN blue edition. (H) Quantification of the normalized intensity mean value of SHH (Alexa 647) staining of the RGCs captured with the same laser intensity from (G). HuFc: N=52, CCN1-HuFc: N=51, P=0.0129. DAPI (blue) was stained for the nucleus. Scale bars: 100 μm for (C), 2 μm for (G). Error bar columns represent mean ± SEM, two-way ANOVA was used for statistical test in (D and F), unpaired student’s t test with welch’s correction was used for the statistical test in (G), *P<0.05; ** P < 0.01; *** P < 0.001; **** P < 0.0001.

### Heparin-binding is required for CCN1 to maintain neural stem cells

Previous studies have identified the two heparin-binding motifs in the human CCN1 protein, which are short peptides rich in lysine (K) and arginine (R) (Chen et al., 2000). The motif is generally characterized as XBBXB, where B refers to the basic amino acid, namely K or R, and X refers to any other amino acid (Margalit et al., 1993). Heparin has been reported to be capable of enriching various growth factors, such as FGF2 and SHH, and regulating their downstream signaling activities (Caldwell et al., 2004; Caldwell and Svendsen, 1998; Conrad, 1997; Williams et al., 2010). Given that the Shh signaling pathway was disturbed by *Ccn1* conditional knockout or overexpression, we tested the hypothesis that heparin-binding is involved in CCN1 function. We constructed a heparin-binding deficient mutant form of CCN1 (CCN1-DM), of which conserved K or R of mouse CCN1 at the indicated sites were all mutated into Alanine (A) (double mutant) (Figure 6A). Appling the heparin resin that could enrich the heparin-binding proteins, we verified that the interaction between CCN1-DM and heparin was largely impaired compared to that between CCN1 and heparin (Figure 6B). We further analyzed the distribution of GFP+ cells in VZ/SVZ, IZ, and CP of CcKO cortices at E18.5 after IUE with PB-CCN1 or PB-CCN1-DM plasmids at E14.5. We found that PB-CCN1 but not PB-CCN1-DM rescued the distribution defects observed in CcKO-PB-Ctrl to similar patterns in WT-PB-Ctrl (Figure 6C and 6D, in comparison to Figure 5I, CcKO-PB-CCN1: VZ/SVZ: 34.66% ± 1.99%, IZ: 29.17% ± 2.70%, CP: 36.17% ± 2.46%; CcKO-PB-CCN1-DM: VZ/SVZ: 25.87% ± 1.12%, IZ: 21.14% ± 2.76%, CP: 52.99% ± 3.62%), suggesting CCN1’s function is dependent on heparin-binding.

### GPC4-CCN1-heparin complex cooperate to regulate activities of the Shh signaling pathway

Interestingly, IP of CCN1-DM-HuFc with GPC4-CHA showed that heparin-binding-deficient CCN1-DM was able to bind to GPC4 (Figure S6A). To further mapping the GPC4-binding domains in CCN1, we constructed plasmids express IGFBP-HuFc, VWC-HuFc, TSP1-HuFc, and CT-HuFc, CCN1_294_-HuFc (aa. 1-294 of CCN1), CCN1_334_-HuFc (aa. 1-334 of CCN1), and CCN1_δ278-334_-HuFc (aa. 278-334 of CCN1 protein was deleted) (Figure S6B and S6D). Co-IP of GPC4-CHA with the various HuFc tagged forms of CCN1 domains or protein mutants showed that CCN1 interacted with GPC4 through its CT domain (Figure S6C) and that there were multiple interaction sites of GPC4 across the CT domain (Figure S6E and S6F). Although GAS1 was identified in our pull-down assay applying CCN1-HuFc recombinant protein, we detected no direct binding of heparin with GPC4 or GAS1 through heparin resin enrichment analysis (Figure S6G). Our results suggest that although the two heparin-binding sites are either on or neighboring to the CT domain, heparin, CCN1, and GPC4 could form a complex in a non-competitive way.

To figure out how CCN1 and heparin work together, we employed RT-qPCR to test the relative expression level of *Ccn1* to reveal the associations between CCN1 and heparin and analyzed the relative expression level of *Gas1* to indicate the activity of the Shh signaling. We dissociated E13.5 cerebral cortices and cultured 24h *in vitro*, then added combinations of different concentrations of CCN1-HuFc (ranging from 0∼250 ng/ml) and heparin (ranging from 0∼2000 ng/ml) for 12h. We harvested the cells and performed RT-qPCR analysis of *Gas1* and *Ccn1* (Figure 6E). Consistent with the RNA-seq result, We found that the addition of CCN1 at a concentration of 2.5 ng/ml was already sufficient to reduce the relative expression level of *Gas1*. We also found that the addition of heparin alone inhibited Gas1 expression. With the increasing concentration of heparin (200ng/ml and 2000ng/ml), both the inhibitory effect of CCN1 and the expression of *Ccn1* were dramatically increased (Figure 6F). These results implied that CCN1 and heparin work cooperatively to regulate the expression of *Gas1*, which antagonizes the activity of the Shh signaling pathway.

Moreover, to test the hypothesis that CCN1 modulates the accessibility of SHH to RGCs, we co-stained HuIgG and SHH on cells dissociated from E13.5 cortices and incubated 1h with SHH and HuFc or CCN1-HuFc, respectively, after culturing 24h. RGCs with CCN1-HuFc incubation showed higher SHH intensity than that with HuFc incubation (Figure 6G-H, 9.01 ± 0.41 for HuFc, 10.43 ± 0.39 for CCN1-HuFc). We discriminated RGCs from neurons through the size of their nuclei and cell morphology in phase, as the diameters of the nuclei of NES+ RGCs were significantly smaller than that of the TUJ1+ neurons (Figure S6H and S6I, 6.31 ± 0.16 μm for NES+ RGCs, 7.16 ± 0.11 μm for TUJ1+ neurons).

## Discussion

In this study, we first revealed the specific expression and binding cell type of CCN1 during mouse cortical development. We dissected that CCN1 protein functions as an indispensable mediator forming complexes with both heparin and RGC membrane-anchored GPC4 to fine-tuning the activities of NSC-maintaining niche factor signaling pathways, resulting in maintenance of NSC homeostasis during development. Moreover, to summarize our findings with previous discoveries, we proposed the working model for the GPC4-CCN1-heparin complex and their reported interacting NSC-maintaining niche factors, taking the most notably regulated Shh signaling pathway as a downstream example (Figure 7A).

**Figure 7.**
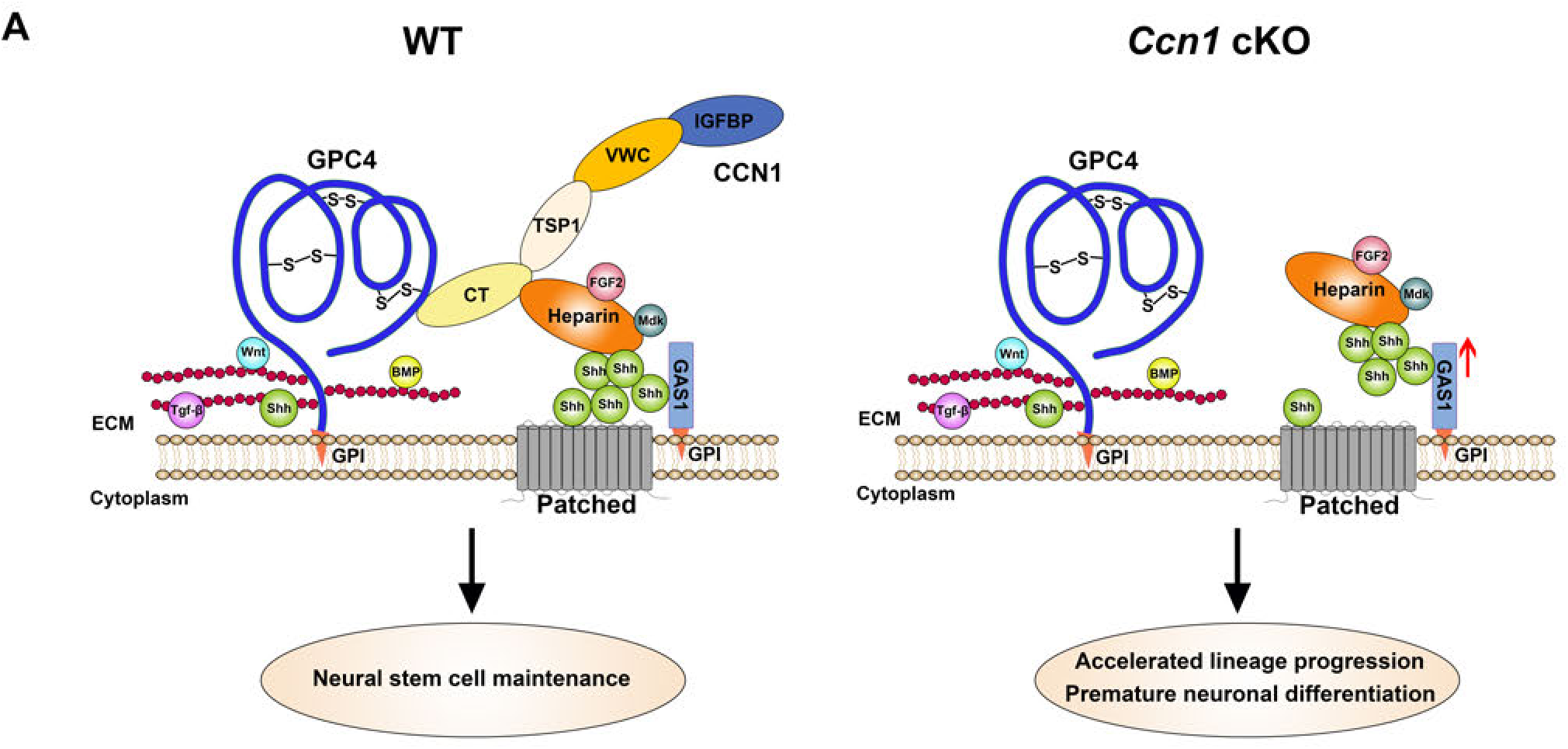
GPC4-CCN1-heparin complex mediates neural stem cell maintenance through regulating niche factor signaling pathway activities. Working model of the GPC4-CCN1-heparin complex during cortical development. In WT mice, CCN1 interacts with GPC4 core protein and heparin through the CT domain, forming a fine-tuning complex for signaling activity regulation. In one aspect, GPC4 anchors on the cell membrane through a GPI anchor, its HS-GAG chain binds with various morphogens, such as WNT, SHH, BMP, Tgf-β. GPC4 helps gradient formation and activity modulation of these morphogens. In the other aspect, heparin can enrich various niche factors, such as SHH, FGF2, Midkine (MDK), and help to regulate niche factor signaling activities to maintain the NSCs. However, in *Ccn1* conditional knockout mice, GPC4 cannot bridge together with heparin without CCN1, interfering with the accessibility of heparin and its enriched niche factors to the cell membrane, resulting in disruption of niche factor signaling activities. For example, the lack of CCN1 upregulates the expression of *Gas1*, which encodes both an Shh co-receptor and an antagonist for the Shh signal activity, leading to a decrease of the Shh signaling activity. The accumulated readout of activity changes of heparin-binding niche factors will eventually accelerate the neural stem cell lineage progression and cause consumptive premature neuronal differentiation.

### The insufficient phenotype in the early corticogenesis

In this study, we applied *Nestin*-Cre (Jackson Laboratory, 003771) mice to induce the conditional knockout of *Ccn1* in the nervous system. However, our results suggested an insufficient knockout efficiency until E15.5∼E16.5 (Figure S2A and S2B) at the end of the peak neurogenesis. Although we detected mild RGC pool size and proliferation deficiency at early embryonic development, the RGC maintenance and the perinatal gliogenesis were severely impaired (Figure 3B-E and Figure 5H-I). These results proved that *Ccn1* was essential for NSC maintenance in late cortical development. As we found that *Ccn1*’s expression is increasing during cortical development, the earlier sufficient knockout of *Ccn1* may indicate a more clear-cut phenotype and interfere with the neuronal differentiation earlier than the summit of the cortical neurogenesis.

### The comprehensive molecular mechanisms of CCN1 protein in maintaining cortical NSCs

From the results that CCN1 distribute and bind on the membrane of RGCs in a clustered punctate manner *in vitro* and *ex vivo* (Figure 1D, 1E, and S1C), we speculated the clustering of CCN1 proteins on the RGC membrane might be vital for their physiological functions. Reminiscent of the clustering of Wnt8 ligands on the cell membrane and their distinctive interaction with N-Sulfo-rich HS clusters rather than N-acetal-rich HS clusters of GPC4, this interaction contributes to the Wnt morphogen gradient formation as well as Wnt signaling activity modulation (Mii and Takada, 2020; Mii et al., 2017). This phenomenon sheds light on the underlying molecular mechanism that CCN1 applies with GPC4. Furthermore, the physiological implications of the clustering of ligands, receptors, and co-receptors may lead to the understanding of the kinetics of niche components.

Furtherly, we selected another heparin-binding growth factor from pulldown-MS data, namely Midkine (MDK), which has been reported to be localized on the basal processes of RGCs and promote the proliferation and self-renewal of stem cells (Xu et al., 2014; Zou et al., 2006). We confirmed the interaction between CCN1 and MDK through IP (Figure S6J), which further supports the hypothesis that the GPC4-CCN1-heparin complex fine-tunes a number of the niche factors and their activities. However, our results suggested that Shh signaling activity regulation might play a core role in maintaining cortical NSCs, which is consistent with the previous understanding that the modulation of Shh activities instead of its complete deactivation plays a vital role in corticogenesis (Yabut and Pleasure, 2018). Whether knockdown of *Gas1* could rescue the premature neuronal differentiation and NSC maintenance defects caused by CCN1 conditional knockout needs further exploration. The mechanisms and functions of how *Gas1* is involved in cortical NSC development will further contribute to the understanding of the NSC behaviors within context-dependent combinations of various niche factor gradients.

Nevertheless, in endothelial cells and fibroblasts, CCN1 was unveiled to interact with lots of integrin receptors. For example, the VWC domain binds with integrin αvβ3 to regulate cell adhesion, proliferation, and apoptosis (Lau, 2011). Integrin αvβ3 activation facilitates the proliferation of both apical and basal NPCs (Stenzel et al., 2014). In the adult V-SVZ niche, CCN1 interacts with integrin α6β1 and regulates EGF signaling pathways, resulting in NSC pool size restriction (Wu et al., 2020). However, integrin family proteins have not been identified in our pulldown-MS data. We ascribe this to the low expression level of integrins during development and the relatively weak interactions between CCN1 and integrins. In other words, the GPC4-CCN1-heparin complex might be the central molecular machinery that is applied by CCN1 to execute its functions during cortical development.

### The physiological significance of GPC4-CCN1-heparin complex formation

Our results further rose the questions about how does the GPC4-CCN1-heparin complex work, and what role does each component play.

GPCs help to shape morphogen gradient through its HS chains and to regulate morphogen or growth factor (such as SHH and FGF2) signaling pathway activities via core protein (Hagihara et al., 2000; Williams et al., 2010; Yan and Lin, 2009). In the Drosophila wing disc, the GPC family homologous Dally and Dally-like proteins (Dlp) help to shape the morphogen gradient through HS side chains, assuring the long-range morphogen ligand concentration around the extracellular membrane (Yan and Lin, 2009). The structural and functional conserved homolog of GPC4 is Dlp, whose membrane-anchored core protein (aa 206-616), rather than Dally, was critical for modulation of activities of the Shh signaling pathway (Williams et al., 2010). We found that GPC4 maintains NSCs depending on the presence of CCN1 (Figure 5H and 5I), and the RNA-seq data suggest more versatile signaling pathways and interacting proteins, including CCN1, are involved in GPC4’s physiological processes (Figure 5F and 5G). However, whether GPC4 is required for NSC maintenance and its underlying pathway activity modulation remains unclear. Our results revealed multiple GPC4 binding sites on the CT domain of CCN1 (Figure S6D-F). Thus, exploring the cortical phenotype of CCN1 binding deficient GPC4 or GPC4 conditional knockout cerebral cortex will illustrate whether GPC4 is necessary for NSCs maintenance. Intriguingly, the different behaviors of GPC4 clusters with N-acetyl-rich or N-sulfo-rich HS chains provide possibilities to fine-tune multiple niche factors with different mechanisms through the GPC4-CCN1-heparin complex (Mii et al., 2017).

Heparin seems to play a role in directly enriching and stabilizing the niche factor ligands, such as FGF2, SHH, and MDK (Figure 7A), hence, mediating the corresponding signaling pathway activity. In drosophila, heparin enhances the interactions of ShhN (19KD Shh N-terminal domain) and its co-receptor iHog, while Dlp cannot interact with Shh ligand directly (Williams et al., 2010). We verified that heparin could not interact with GPC4 or mammalian Shh co-receptor as well as antagonist GAS1 directly (Figure S6G). Consistent with the disrupted NSC-maintaining niche factor signaling pathways after *Ccn1* cKO or overexpression, heparin plays the role of executing the direct ligands enrichment and modulating the ligand-co-receptor interacting strength, which coincides with the necessity of heparin-binding for CCN1 to maintain NSCs.

There are some co-receptors of niche factors, such as SHH, NOTCH, and EGF, being found in CCN1 interacting proteome list (Datasets S2). Whether those co-receptors interact with CCN1 directly or indirectly through GPC4 or heparin and impact the corresponding signaling activity are worthy of exploration. On the fact that membrane anchorage is indispensable for GPC4 to regulate the activity of the Shh signaling (Williams et al., 2010), the physiological significance of CCN1 is potentially to be an adaptor that pulls together the ligands that heparin enriched and the receptors or co-receptors that GPC4 interact with, ensuring the long-range morphogen signaling activities. Further investigations on those hypotheses could provide us in-depth insights into the physiological implications of CCN1.

Intriguingly, ShhN shares a “hotspot” interacting interface for its receptors, such as Cdo and Gas1, as well as heparin and Chondroitin Sulfate (CS) side chains of HSPGs (Whalen et al., 2013). Thus, the GPC4-CCN1-heparin complex provides access for positive and negative regulators, including niche factors and their corresponding receptors and co-receptors, to NSCs simultaneously. It works as a modulator to integrate and fine-tune the final signal input of niche factors for NSCs. Finally, different gradient combinations of these regulators will lead to a distinct phenotypic output of NSCs.

### Context-dependent multifaceted physiological functions of CCN1

Interestingly, GPC4 was also identified in our adult V-SVZ niche pulldown by CCN1-HuFc, albeit with a relatively large lower enrichment ranking (Wu et al., 2020), which means the GPC4-CCN1-heparin complex might also contribute to the functionality of CCN1 within the adult NSC niche. However, not only the abundance of the interacting membrane proteins but the phenotypic readout of CCN1 during embryonic cortical development differs from that of the adult V-SVZ NSCs. Moreover, the expression pattern of *Ccn1* and *Gpc4* in the germinal zone of pallium and subpallium is also different during development (Figure 1A and 4F). Prominently, *Ccn1* shows a higher expression abundance in subpallium than pallium, while *Gpc4* expresses higher in cortex and LGE than MGE. Combining with the multitudinous niche factors, such as SHH and WNT, with gradient formation in a dorsal-ventral or anterior-posterior axis, the behaviors of NSCs localized in the brain subdivisions could be different. Indeed, the proliferative potentials, the persistence, and the diversifications of the specified progenies of NSCs in the dorsal cortex, in subpallial LGE, or MGE at the same stage are quite distinctive. Subpopulations of LGE NSCs are preserved to be the adult V-SVZ NSCs. However, the cortical neurogenic NSCs exhaust perinatally. While MGE NSCs proliferate and generate interneurons instead of pyramidal projection neurons in earlier gestational age (Hu et al., 2017; Miyoshi et al., 2007).

These lines of evidence potentially suggest different functions of CCN1 in subpallial NSCs and imply a dynamic context-dependent multifaceted physiological processes of CCN1. The corresponding NSC niche components that contribute to the maintenance, proliferation, differentiation, and fate determination of NSC, satisfied the developmental and aging needs of NSCs within subregions of the brain.

## Supporting information

Key Resource Table

## AUTHOR CONTRIBUTIONS

J-L.Z. designed and conducted experiments, performed data analysis, and wrote the manuscript; S-J.Q. designed and conducted experiments, H.W. bred mice, conducted experiments and data analysis; J.W. bred mice and helped to discuss the project; G.C., H-H.J.W., and H.P. helped construct plasmids; J.L. and C-Q.C. helped with experiments; K.T. helped generate the volcano plot. Q.S. supervised the project and revised the manuscript.

## DATA AVAILABILITY

Raw fastq sequence data from the Illumina NovaSeq platform and gene expression matrices are available on National Center for Biotechnology Information Gene Expression Omnibus (Accession No. GSE165861).

## ACKNOWLEDGMENTS

We thank Dr. Lester F. Lau for providing *Ccn1*^flox/flox^ mice, Dr. Peng Yang from Tongji University for providing suggestions on CCN1 mutation, Dr. Xiaohua Shen from Tsinghua University for providing *PiggyBac*-pCAG-IRES -EGFP plasmid, Dr. Xiao-ling Hu from Capital Medical University for guidance on acute slice culture, ZEISS Microscopy Customer Center Shanghai for providing professional service of Axio Scan.Z1 Slide Scanner System. This study was supported by grants from NSFC (31200798, 20131351353), Tsinghua CLS, and Tongji University.

## STAR METHODS

Detailed methods are provided in the online version of this paper and include the following:

### Contact for Reagent and Resource Sharing

Further information and requests for resources and reagents should be directed to and will be fulfilled by the Lead Contact, Qin Shen

### Experimental Model and Subject Details

#### Animals and generation of *Ccn1* conditional knockout mice

Mice were bred and maintained in the animal facility at the Center of Biomedical Analysis in Tsinghua University and the animal facility of Tongji University. All animal protocols used in this study were approved by the IACUC (Institutional Animal Care and Use Committee) of Tsinghua University and Tongji University and performed following the guidelines of the IACUC. The laboratory animal facility has been accredited by AAALAC (Association for Assessment and Accreditation of Laboratory Animal Care International). ICR mice were obtained from Vital River Laboratory Animal Technology Company (Beijing, China). Neural-specific *Ccn1* conditional knockout mice were generated by breeding *Nestin*-Cre (B6.Cg-TgN(Nes-cre)1Kln)/J, Stock No: 003771) with *Ccn1*^flox/flox^ mice, which were a gift from Prof. Lester F. Lau (Kim et al., 2013). In all experiments, mice with genotypes of *Ccn1*^fl/fl^ mice were used as the control mice. Mid-day of the vaginal plug identified was calculated as embryonic (E) day 0.5.

#### Primary neural stem cell culture

Embryos from sacrificed timed-pregnant mice were microdissected under a stereomicroscope using fine tweezers. For electroporated brains, the EGFP positive cortices were dissociated with papain. The dissociated cells were cultured 48h on poly-L lysine (PLL) coated coverslip before immunostaining. For the heparin and CCN1-HuFc protein functional test, one piece of the cortex for each PLL coated well of 24-well plate was prepared, twelve E13.5 brains were dissociated independently as one individual repeat. The dissociated cells were plated and cultured 24h before 12h treatment with orthogonal combinations of heparin (0, 0.2, 2, 20, 200, 2000 ng/ml) and CCN1-HuFc protein (0, 2.5, 25, 250 ng/ml) concentration. For SHH with HuFc or CCN1-HuFc protein binding assay, cells were dissociated from E13.5 cortices and cultured 24h before 1h treatment with 50 ng/ml SHH combined with 20ng/ml HuFc or 25ng/ml CCN1-HuFc proteins, respectively. The NSC culture conditions and medium were following the adherent NSC culture procedures as described previously (Qian et al., 2000).

#### Acute slice culture

Pregnant mouse with E13.5 embryos was anesthetized by intra-peritoneally injection of 1% (W/V) pentasorbital sodium. The uterine horns were pulled out from abdominal cavity using forceps and scissors, embryonic brains were exposed by fine tweezers and quickly embedded with 3% LMP (low melting) Agarose (Genview) in Tissue-Tek Cryomold (Sakura). 200 μm coronal sections of forebrain were made in cold hibernation solution (30mM KCl, 5mM NaOH, 5mM NaH_2_PO_4_, 0.5mM MgCl_2_, 20mM Sodium Pyruvate, 5.5mM Glucose, 200mM Sorbitol, pH7.3-7.4) using vibratome. Then the slices were incubated in NSC medium (DMEM with 0.16 g/L NAC, N2/B27 supplement, 10 ng/ml bFGF) with 12.5 ng/μl HuFc or 25 ng/μl CCN1-HuFc protein at 37℃ for 30min. The slices were fixed by 4% paraformaldehyde (PFA) in phosphate buffer (PB) (short for “PFA” thereafter) 2-4h, and immunostained following the immunostaining procedures, specially, anti-Human IgG1 Fc, Alexa Fluor 488 secondary antibody was used to recognize HuFc peptide.

#### *In utero* electroporation (IUE)

Timed-pregnant mice were anesthetized by isoflurane, putting on a heated pad. The uterine horns were exposed, and 1 μl of plasmid mixture with fast green dye (p-base : *PiggyBac*: 3:1 ∼2 µg/µl) was injected into the lateral ventricle of the brain with a 5-gauge microinjection needle. Embryos were then clamped between homemade 10 mm-diameter tweezers-type disc electrodes. Five 50 ms on, 1s off electrical pulses were applied to each embryos using an electroporator (ECM830, Harvard Apparatus). 28V, 32V, or 36 V were applied to E12.5, E13.5, or E14.5 embryos correspondingly. Embryos were then placed back into the abdominal cavity to continue normal development.

#### Bromodeoxyuridine (BrdU) labeling and detection

BrdU (dissolved in saline) was injected intra-peritoneally into E14.5 pregnant mice proportional to body weight, 100mg/Kg accordingly. Mice were sacrificed 24h after BrdU injection for cell cycle exit index analysis. The frozen sections were pretreated with 500U/ml DNase-I dissolved in DNase-I buffer (40 mM Tris–HCl, 10 mM NaCl, 6 mM MgCl_2_, 10 mM CaCl_2_, pH 7.9) 10min at RT and washed by PBS 3 times. Then, the frozen sections were processed following the immunostaining protocol using BrdU and Ki67 antibodies.

### Method Details

#### Cell line culture and transfection

293FT and Neuro2a (N2a) cell lines were cultured by DMEM (Corning) with 10% Fetal Bovine Serum (FBS) and Penicillin-Streptomycin (P/S). For the following IP or heparin resin enrichment, the plasmids were co-transfected into 293FT cells with 3 μg plasmids (1.5 μg each plasmid for Co-IP) in total by Exfect (Vazyme Biotech, plasmids: Exfect=1:1.5). For the following immunostaining, the plasmids were co-transfected into N2a cells with 1.5 μg each plasmid by Exfect (plasmids: Exfect=1:2).

#### Brain frozen section and immunostaining

Embryonic brains of the indicated stage were fixed 8-16h accordingly in PFA, while the postnatal puppies were perfused and post-fixed overnight by PFA. The fixed brains were dehydrated by 30% sucrose in PBS until sinking and embedded using O.C.T. (Sakura). Then 20 μm serial coronal sections were made for embryonic brains with six slides intervals, while postnatal brains with 30 μm thickness with eight slides intervals. For nuclear marker staining, the slices were pre-treated with 10 μg/ml proteinase K for 10 min at RT. After washing, the slides were blocked in 5% BAS with 0.3% Triton-X 100 (short for blocking buffer later) and then incubated with the primary antibodies with proper dilution in blocking buffer at 4℃ overnight, washed with PBS for three times and proceeded to the secondary antibody and DAPI incubation for 1h at RT. The slides were mounted by fluoromount and coverslip.

The primary antibodies used in this paper are included in the Key Resources Table.

#### Protein lysate pulldown

Ten E13.5 forebrains were dissected in hibernation solution and cut into pieces before being lysed in 1ml IP lysis buffer (25 mM Tris•HCl pH 7.4, 150 mM NaCl, 1% NP-40, 1 mM EDTA, 5% glycerol). The supernatant after centrifuge was aliquoted into two tubes and incubated with 20 μl Dynabeads protein G and 1.25 μg HuFc or 2.5 μg CCN1-HuFc recombinant protein, respectively, in 4℃ overnight. The supernatants were discarded after beads enrichment through the magnetic frame. Then the beads were washed by IP lysis buffer on the rotator. The eluate was collected for mass spectrometry detection.

#### Construction of plasmids

A multiple cloning site (MCS) was inserted into pFUGW-H1-EGFP/RFP vector to construct pUbC-MCS-EGFP/RFP, the CDS sequence without stop codon of *Ccn1* or *Gpc4* was inserted into the MCS for CCN1-EGFP/RFP or GPC4-EGFP fusion protein expression. The DNA fragments of CCN1_294_, CCN1_334_, CCN1_δ278-334_, and CCN1 without stop codon were inserted into pCAG-HuFc (human IgG1 Fc). The CDS sequences of GPC1/2/4/6, SDC1/2/3/4, GAS1, MDK, GPC4 δHS, and GPC4 core proteins used for Co-IP were inserted into pUbC-CHA-IRES-EGFP vectors through restriction enzyme digestion and T4 ligase ligation. The site mutations of heparin-binding deficient CCN1-DM and GPC4 δHS and the deletions for CCN1_δ278-334_ were all prepared using site-directed overlap PCR. *PiggyBac*-pCAG-IRES-EGFP (PB-Ctrl) were constructed as described previously (Hu et al., 2017). *PiggyBac*-pCAG-CCN1-IRES-EGFP (PB-CCN1), *PiggyBac*-pCAG-CCN1-DM-IRES-EGFP (PB-CCN1-DM), and *PiggyBac*-pCAG-GPC4-IRES-EGFP (PB-GPC4) were constructed by inserting the CDS sequences of CCN1, CCN1-DM, or GPC4 fragments into PB-Ctrl vector using Pro Ligation-Free Cloning Kit.

Mouse cDNA was used as the template to amplify desired DNA fragments by PCR for cloning. DNA sequencing was performed to confirm all the desired mutations and in-frame fusions. All oligonucleotides used for the PCR amplification were included in the Key Resources Table.

#### Immunoprecipitation (IP) and heparin resin enrichment

293FT cells were lysed 48h after transfection using 200 μl IP lysis (25 mM Tris•HCl pH 7.4, 150 mM NaCl, 1% NP-40, 1 mM EDTA, 5% glycerol) for 2-4h at 4℃. An equal volume of the supernatant after centrifugation was denatured for each sample as the input group. While the remaining lysates were incubated with Dynabeads Protein G (with high affinity to HuFc peptide) or heparin resin at 4℃ for 6-12h, the Dynabeads were washed by IP lysis on magnetic frames for three times, while the heparin resin was washed through centrifugation. The elutes from beads (or resin) saved as the IP (or the enriched group) were resuspended and denatured for immunoblotting. Each IP and heparin resin enrichment experiment was performed at least three times.

#### Western blot

For IP, 15 μl of input or IP were applied in each lane. For heparin resin enriched samples, 5 μl of input and 15 μl of the enriched sample were loaded in each lane. The immunoblotting was performed using mouse anti-HA primary antibody, the secondary HRP conjugated antibodies anti-mouse or anti-human IgG was added together or separately if the molecular weight of the target proteins is similar. For CCN1 overexpression efficiency detection, CCN1 protein was detected firstly, then β-Actin antibody was applied to the same PVDF membrane after stripping 15 min at room temperature (RT).

#### RNA extraction and Real time-quantitative PCR (RT-qPCR)

Total mRNA extraction was performed following the TRIzol^®^ reagent protocol for both RT-qPCR and bulk RNA-seq. Before RT-qPCR, the cDNA library was reverse transcribed using random primers after gDNA removal following the manufacturer’s protocols of One-step gDNA Removal and cDNA Synthesis SuperMix Kit. qPCR reactions were set up using the AceQ qPCR SYBR Green Master Mix (Transgene). Reactions were run on a Bio-Rad CFX Real-time 384X PCR system. Primers for *Gapdh*, *Ccn1*, *Gas1* have been included in Key Resources Table.

#### NGS mRNA library preparation

500-1000ng total RNA was prepared for mRNA library preparation, the mRNA library preparation including mRNA capturing, fragmentation (200-300bp), adaptor (NEB, Ultra II RNA Library Prep Kit for Illumina) ligation, and library amplification and purification was finished following the manufacturer’s protocols of KAPA Hyper Prep Kit (Kapa Biosystems).

#### Bioinformatic analysis of Illumina mRNA NGS data

Clean Data from the Illumina NovaSeq PE150 RNA sequencing were uploaded to the online website https://usegalaxy.org/ for bioinformatic analysis. The quality control for clean data was processed by Trimmomatic R package, parameters including LEADING 20, TRAILING 20, and MINLEN 35. The data were mapped to mm10 genome with default parameters using HISAT2, the output BAM file was calculated to counts including only primary counts with featureCounts R package, and the fold change and P-value of DEGs were calculated by DESeq2. FPKM were calculated using counts to FPKM formula in R software. The volcano plot was generated with the ggplot2 R package. GO enrichment analyses were processed on the online website http://metascape.org. GSEA was performed with fold change-ranked gene counts data by GSEA software downloaded from http://software.broadinstitute.org/gsea/downloads.jsp. The enrichment results were visualized with Graphpad Prism 6.

#### *In situ* hybridization

The brain sections were fixed by 4% PFA at RT for 20min and washed twice with PBS. Then the sections were treated with 10 μg/ml proteinase K (Roche) dissolved in proteinase K buffer (5mM EDTA, 50mM Tris-HCl, pH7.4∼7.5) for 10min. Next, fix the sections with 4% PFA again for 10min and rinse with PBS. Subsequently, acetylate the sections in dark in acetylation solution (1.3% triethanolamine, 0.25% acetic anhydride, 17.5mM HCl). Rinse the sections twice and pre-hybridized in hybridization buffer (50% formamide, 250ug/ml yeast RNA, 500ug/ml herring sperm DNA, 5X Denhardts solution, 5X SSC, RNase-free, pH7.0) for 1h. Digoxin-labeled RNA probes were incubated at 85℃ for 5min and chilled on ice. Hybridize overnight in 65℃ incubator with 0.1ng/μL RNA probe in hybridization buffer. After rinsing with 0.2X SSC twice at 65℃ for 20min each time, the sections were rinsed with 0.2X SSC at RT again and subsequently rinsed with B1 buffer (0.1M Tris-HCl, pH7.4∼7.5) twice for 5min. Then block the sections with blocking buffer (10% goat serum in B1 buffer) for 1h, and incubate the sections with anti-digoxin antibody (Roche, 1:5000) in blocking buffer at 4℃ overnight. Wash the sections with B1 buffer three times and balance with B2 buffer (0.1M Tris-HCl, pH9.5, 0.1M NaCl, 0.05M MgCl2, 0.1% TWEEN-20) twice. Finally, incubate the sections with 50×NBT/BCIP in B2 buffer in dark, check the slides periodically and terminate the reaction with 10mM Tris (pH 8.0), 0.05M EDTA.

#### Imaging and cell counting

Immunofluorescent slides were imaged by Zeiss LSM 880 fluorescent inverted microscopy, images in Figure 1D, 2F, 3H, 5A, and S5E were captured with stacks of 10 slices with 1.4 μm Z step size and shown after maximum intensity projection, images in Figure 1B, 2A, 2C, 2H, 3B’’, 4G, 5H, 6C, S1C, S2C, S3A, and S4D were snaped with the same optical depth. Images in Figure 3B’ were taken by Zeiss Observer.D1 inverted microscopy, while images in Figure 3F were taken by Zeiss Axio Observer 3 inverted microscopy. Cell counting and regional GFP intensity mean were performed using ZEN blue edition in 1-5 sections with anatomically matched positions in experimental groups. ISH images were taken by Zeiss Axio Scan.Z1. The western blot images were taken by Bio-Rad Imaging System.

#### Quantification and Statistical Analysis

Statistical analyses were performed with Student’s t test with Welch’s correction, one-way ANOVA followed by Dunnett’s posttest, or two-way ANOVA followed by Sidak’s posttest using GraphPad Prism 6 Software. Data are presented as mean ± SEM, p < 0.05 was considered significant. The value of n for each graph was stated in the Figure legend.

## Supplemental Information

### Supplemental Figure Legend

**Figure S1. Related to Figure 1.**
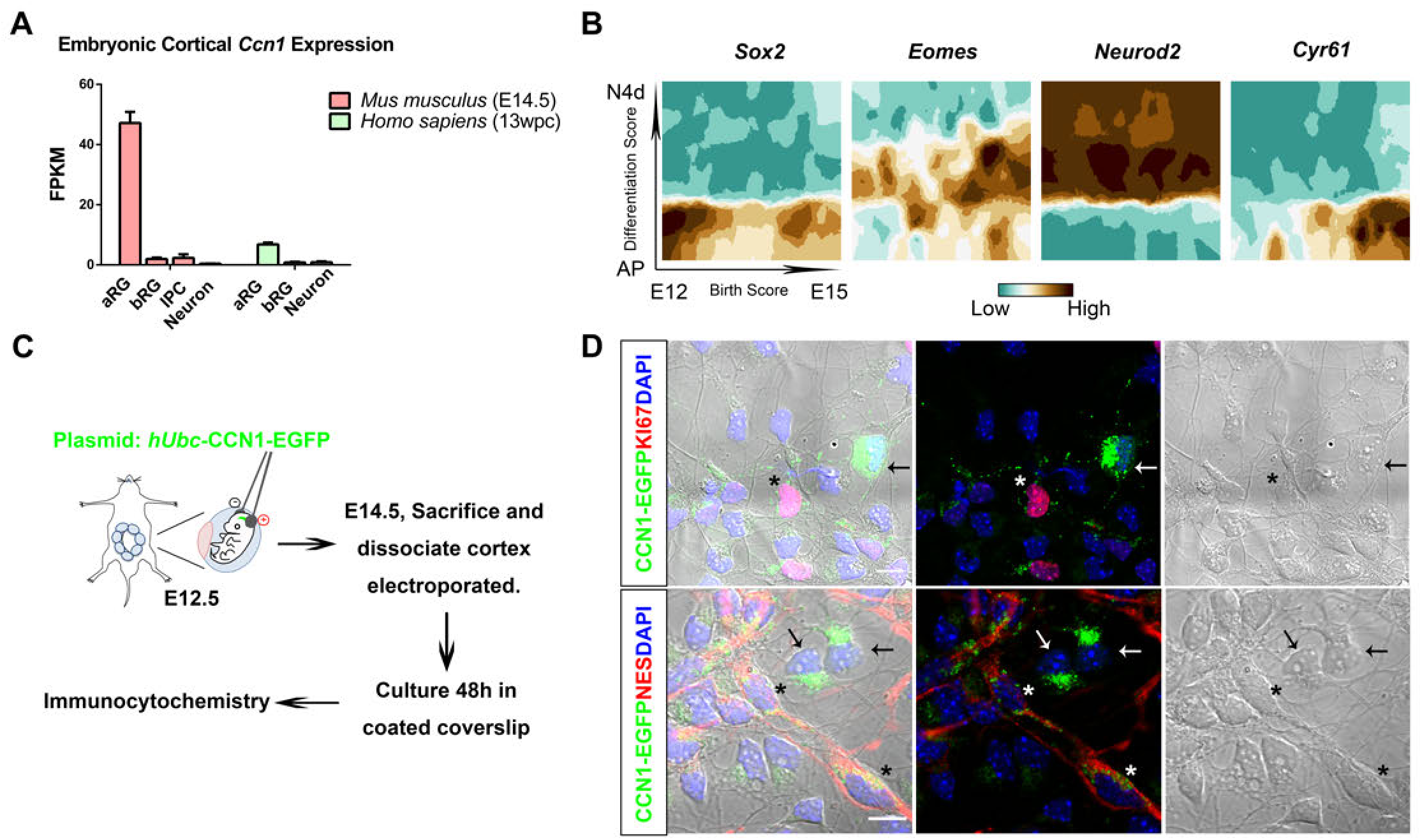
*Ccn1* is increasingly expressed in apical radial glial cells during corticogenesis; CCN1 protein localizes on radial fibres of proliferative NPCs in a clustered punctate manner *in vitro*. (A) Bar graph of the FPKM (fragments per kilobase per million mapped reads) value of *Ccn1* in different purified cell types of E14.5 mouse cortex and 13 weeks postconception (wpc) human cortex. (B) Dynamic expression gradient map of *Sox2*, *Eomes*, *Neurod2*, and *Cyr61* of the birthdate and differentiation status by t-SNE clustering throughout mouse corticogenesis. (C) Schematic diagram of experimental design for immunocytochemistry handled on div2 E14.5 cortical culture on PLL-coated coverslips from dissociated cells after IUE of *hUbC*-CCN1-EGFP (green) plasmid in embryos at E12.5. (D) Representative images of the cells immunostained by KI67 (upper, red) or NES (lower, red), following the procedures from (B). Arrows point to postmitotic neurons; asterisks point to RGCs or proliferating cells. aRG: apical radial glia, bRG: basal radial glia, IPC: intermediate progenitor cell, N: neuron, AP: apical progenitor, N4d: 4-day-old neurons. Scale bar: 10 μm for (D).

**Figure S2. Related to Figure 2.**
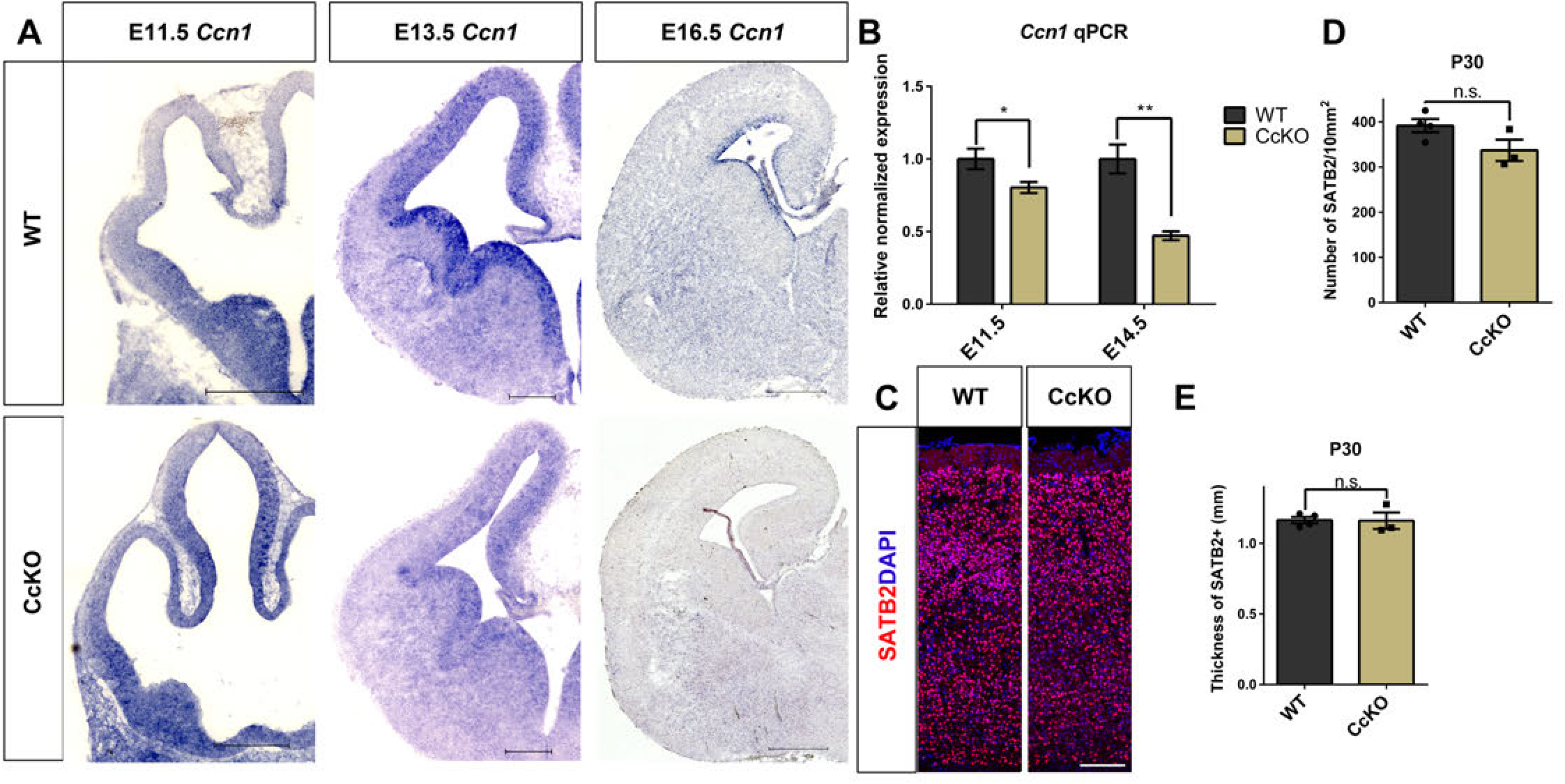
The gradually complete conditional knockout of *Ccn1* does not alter neurogenesis significantly postnatally. (A) *In situ* hybridization of *Ccn1* on the coronal forebrain sections of WT and CcKO mice atE11.5, E13.5, and E16.5. (B) Bar graph of the relative normalized expression of *Ccn1* in the CX of E11.5 and E14.5 brain by RT-qPCR. *Gapdh* was used as the reference gene, and each normalized gene expression was divided by the mean of *Ccn1* expression in WT. WT: N=6, CcKO: N=7, P=0.0422 (E11.5); WT: N=6, CcKO: N=4, P=0.0023 (E14.5). (C) Representative images of SATB2 staining on P30 brain sections of the WT or CcKO cortices. (D-E) Statistical bar graph of the number of SATB2+ projection neurons in every 10 mm^2^ unit area (D), and the thickness of the SATB2+ Layer II-VI in WT and CcKO mice at P0 (E). WT: N=4, CcKO: N=3, P=0.1305 (D), P=0.9553 (E). Scale bar: 500 μm for (A), 200 μm for (C). Error bar columns represent mean ± SEM, unpaired student’s t test with Welch’s correction was used for the statistical test in (B, D, and E), n.s. P>0.05, *P<0.05; ** P < 0.01.

**Figure S3. Related to Figure 3.**
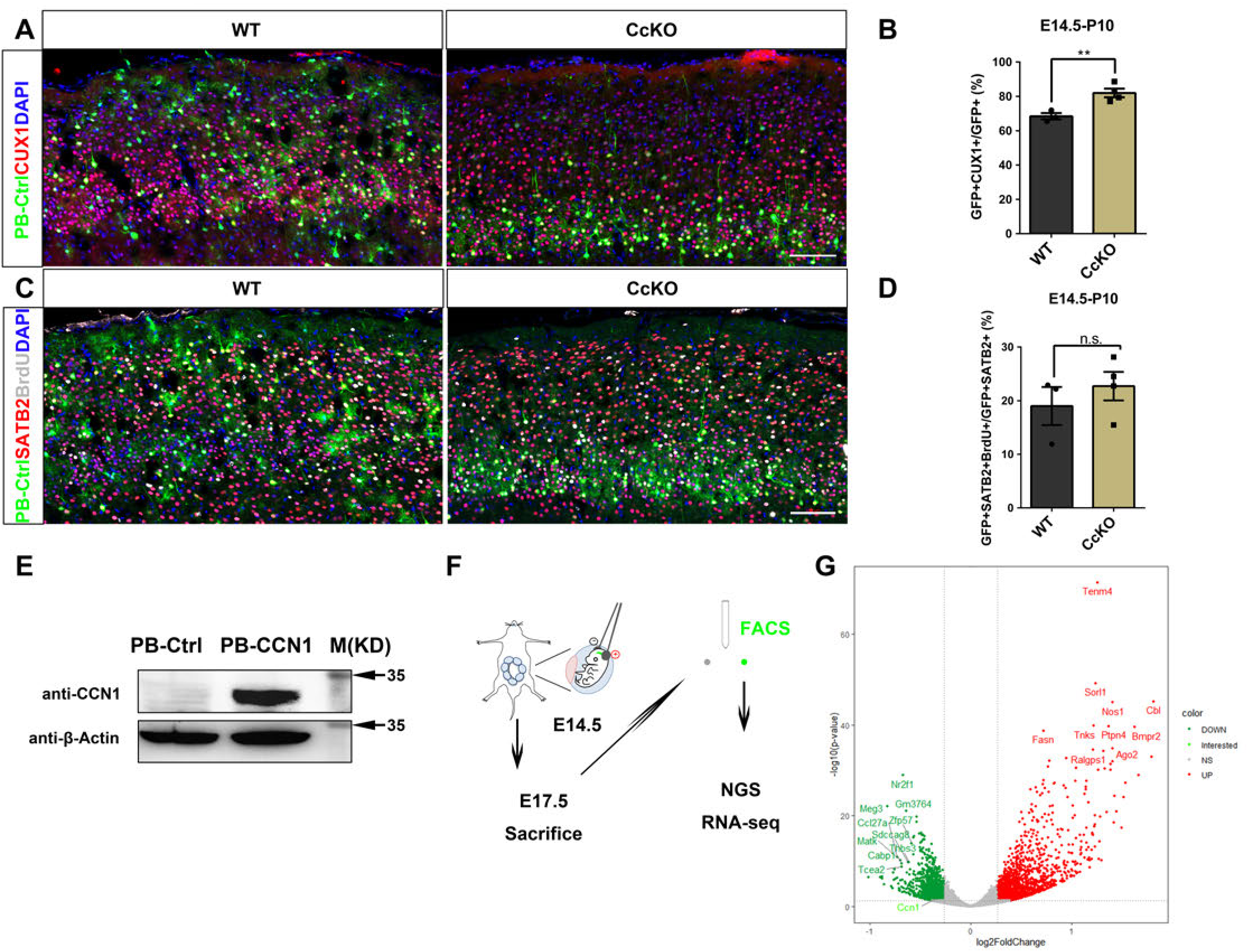
Loss of *Ccn1* reduces gliogenesis, but does not alter rate of neurogenesis significantly at E15.5. (A) Representative images of CUX1 (red) immunostaining in UCP of WT or CcKO brain sections at P10 after IUE of PB-Ctrl (green) at E14.5. (B) Quantification of the percentage of CUX1+ neurons in UCP GFP+ cells in the WT or CcKO mice cortices from (C). WT: N=3, CcKO: N=4, P=0.0074. (C) Representative images of BrdU (white) and SATB2 (red) co-immunostaining in UCP of WT or CcKO brain sections at P10 after IUE of PB-Ctrl at E14.5. (D) Quantification of the percentage of E15.5 generated GFP+SATB2+BrdU+ cells in GFP+SATB2+ neurons from (A). WT: N=3, CcKO: N=4, P=0.4487. (E) Western blot of protein lysates of PB-Ctrl and PB-CCN1, respectively, which were transfected in 293FT cells using CCN1 and β-Actin antibodies. (F) Schematic diagram of the experimental design of IUE (E14.5 → E17.5) with plasmids in pregnant mice with WT and CcKO embryos, purification of GFP+ cells by FACS, and the following RNA-seq analysis. (G) Volcano plot of log2FoldChange (X-axis) and –log10(P-value) (Y-axis) for differentially expressed genes of processed Illumina RNA-seq data of WT and CcKO samples from Figure S3E. The vertical dashed line indicates Fold Change=±1.2, and the horizontal dashed lines indicate P-value=0.05. Red dots indicate significantly up-regulated genes with Fold Change>1.2 and P-value<0.05, dark blue dots indicate significantly down-regulated genes with Fold Change<-1.2 and P-value<0.05, and grey dots indicate not significantly differentially expressed genes with −1.2<Fold Change<1.2 or P-value>0.05. The downregulation of *Ccn1* was highlighted with Fold Change <-1.2. Scale bar: 100 μm for (A) and (C). Error bar columns represent mean ± SEM, unpaired student’s t test with Welch’s correction was used for the statistical test in (B) and (D), P>0.05 for n.s., **P<0.01.

**Figure S4. Related to Figure 4.**
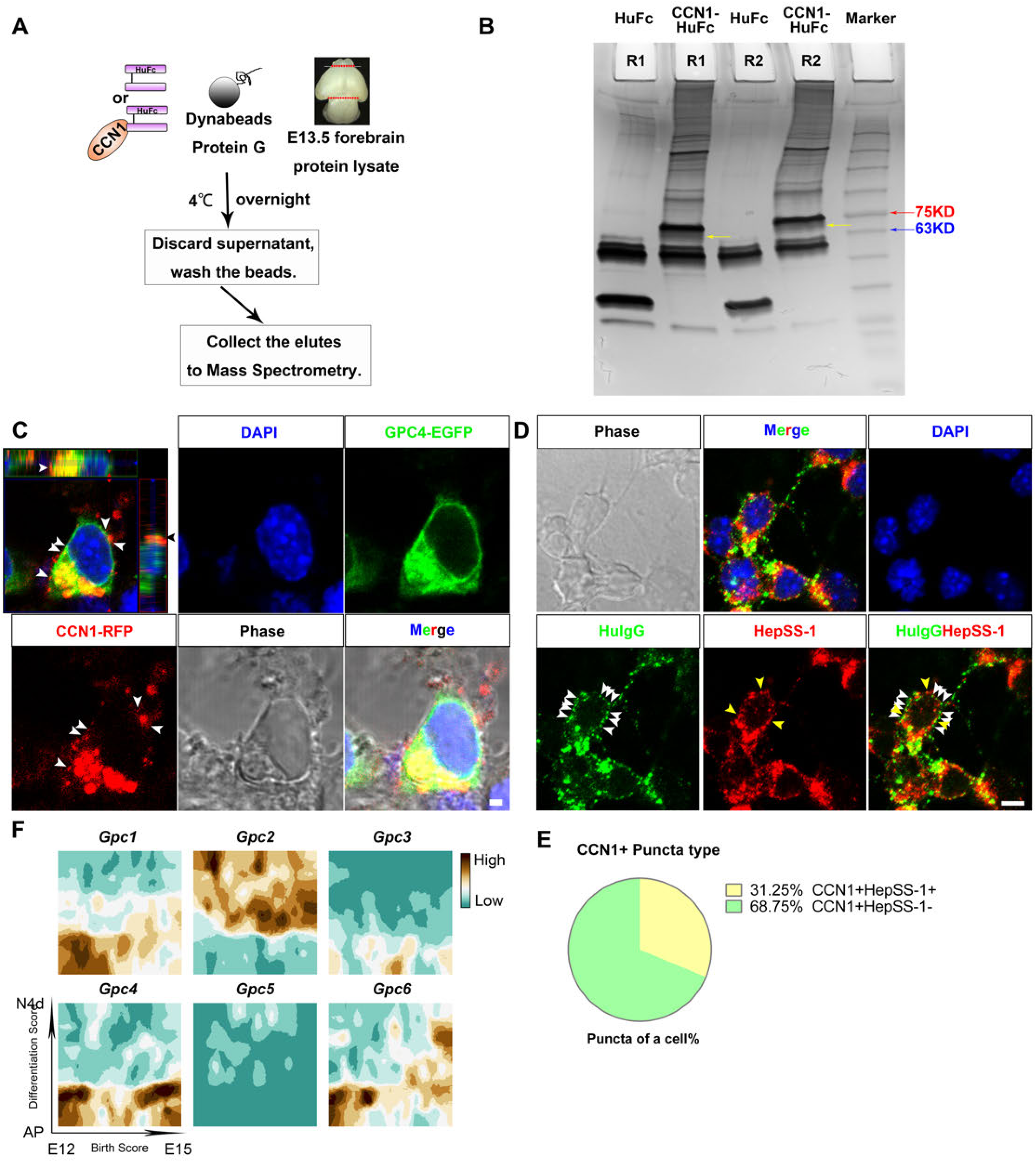
CCN1 interacts with GPC4 on the RGC membrane. (A) Schematic flowchart of HuFc or CCN1-HuFc recombinant bait protein pull-down with E13.5 forebrain protein lysate following mass spectrometry qualitative analysis. (B) Image of silver staining for two repeats of HuFc or CCN1-HuFc pulldown proteins separated through SDS-PAGE. Yellow arrows point to the band of GPC4. (C) Images of N2a cells expressing both GPC4-EGFP and CCN1-RFP fusion proteins 48h after transfection. Arrowheads point to colocalized CCN1 and GPC4 on the cell membrane. (D) Representative images of the co-immunostaining of HuIgG (green) and N-Sulfo HS (HepSS-1, red) on RGCs 24h after dissociated from E13.5 cortices and 0.5h incubation with CCN1-HuFc. White arrowheads point to CCN1+HepSS-1-puncta, yellow arrowheads point to CCN1+ HepSS-1+ puncta on the RGC membrane. (E) Quantification of the percentage of CCN1+HepSS-1- and CCN1+ HepSS-1+ puncta type. N=7 RGCs. (F) Dynamic expression gradient map of *Gpc1, Gpc2*, *Gpc3*, *Gpc4, Gpc5*, and *Gpc6* of the birthdate and differentiation status by t-SNE clustering throughout mouse corticogenesis. DAPI (blue) were stained for nuclei. Scale bar: 2 μm for (C), 5 μm for (D).

**Figure S5. Related to Figure 5.**
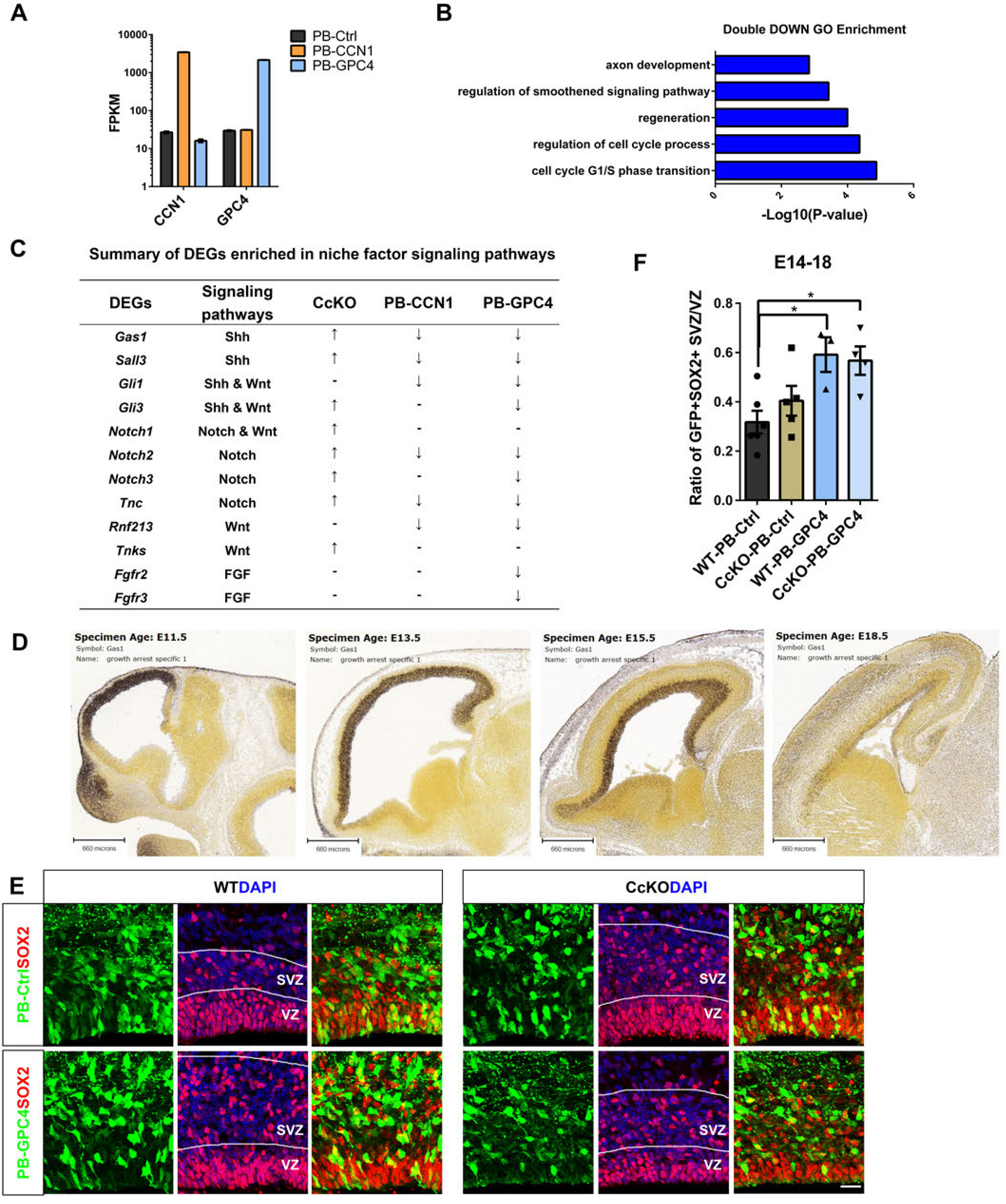
GPC4 and CCN1 regulate multiple common signaling pathways related to NSC maintenance, while GPC4 facilitates dislocated NPCs in SVZ independent of CCN1. (A) Mean FPKM bar graph of *Ccn1* and *Gpc4* from RNA-seq data of PB-Ctrl, PB-CCN1, and PB-GPC4. The Y-axis scale was presented in log10 fashion for better visualization of the reduction of *Ccn1* expression when *Gpc4* was overexpressed. (B) Bar graph for GO enrichment of PB-CCN1 and PB-GPC4 double down-regulated DEGs. (C) Summary chart of the regulatory tendency for significantly DEGs enriched in Shh/Wnt/Notch/FGF signaling pathways from RNA-seq of CcKO, PB-CCN1, and PB-GPC4, respectively, compared to their controls. (D) *In situ* hybridization of *Gas1* during indicated embryonic developmental stages, data are from the *Allen Brain Atlas* database. (E) Representative images for the zoom-in VZ/SVZ region from Figure 5H immunostained by SOX2. Dashed lines indicate the border of VZ and SVZ. (F) Quantification of the ratio of SOX2+GFP+ progenitor cells in SVZ to VZ in the brain sections from (H) immunostained with SOX2 (J). WT-PB-Ctrl: N=3, CcKO-PB-Ctrl: N=3, WT-PB-GPC4: N=3, CcKO-PB-GPC4: N=3. P<0.05 (WT-PB-GPC4 vs. WT-PB-Ctrl, CcKO-PB-GPC4 vs. WT-PB-Ctrl). Scale bar: 660 μm for (D), 20 μm for (E). Error bar columns represent mean ± SEM, one-way ANOVA was used for statistical test in (F). *P<0.05.

**Figure S6. Related to Figure 6.**
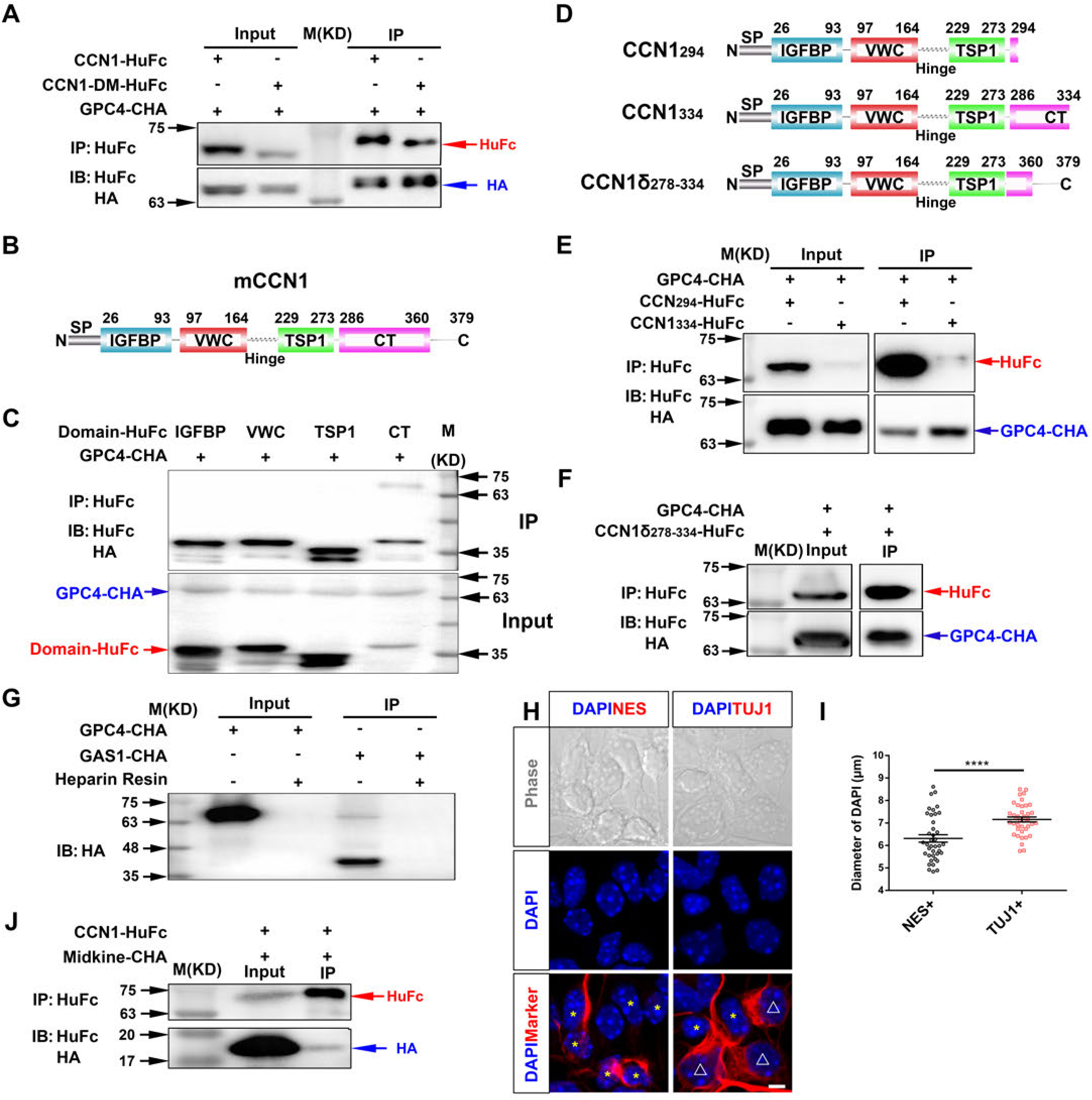
GPC4 has multiple interaction sites on the CT domain of CCN1 and forms a complex with heparin through CCN1 in a non-competitive way. (A) Western blot for input or IP of GPC4-CHA and CCN1-HuFc or CCN1-DM-HuFc, respectively. M: marker. (B) Schematic diagram of the four domains of CCN1 protein. (C) Western blot for input (lower) or IP (upper) of GPC4-CHA and IGFBP-, VWC-, TSP1-, or CT-HuFc, respectively. M: marker. (D) Schematic diagram of the construction of CCN1_294_, CCN1_334,_ and CCN1δ_278-334_ mutants of CCN1 protein. (E) Western blot for input or IP of GPC4-CHA and CCN1_294_-HuFc and CCN1_334_-HuFc, respectively. M: marker. (F) Western blot for input or IP of GPC4-CHA and CCN1δ_278-334_-HuFc. M: marker. (G) Western blot of GPC4-CHA or GAS1-CHA after heparin resin enrichment, lanes without heparin resin are input groups. (H) Representative images of phase, DAPI (blue), and NES (left, red) or TUJ1 (right, red) of cells cultured 24h after dissociation from E13.5 cortices. Yellow asterisks mark the RGCs, white triangles mark the neurons. The corresponding RGCs in the phase were rounded shaped in morphology, while the corresponding neurons were morphologically more flatted and multihole on the surface. (I) Quantification of diameters of the nuclei of NES+ RGCs and TUJ1+ neurons. N=39 for NES+ cells, N=39 for TUJ1+ cells, P<0.0001. (J) Western blot for input or IP of Midkine-CHA and CCN1-HuFc. M: marker. DAPI (blue) were stained for nuclei. Scale bar: 5 μm for (H). Unpaired student’s t test with Welch’s correction was used for (I). ****P<0.0001.

### Supplemental tables

**Table 1. Related to Figure 3**

**E14-17 PB-Ctrl CcKO_WT gene expression & Fold Change**

**Table 2. Related to Figure 4**

**Identified Proteins from CCN1-HuFc or HuFc pulldown**

**Table 3. Related to Figure 5**

**E13-15 PB-Ctrl_PB-CCN1_PB-GPC4 gene expression & Fold Change**

